# Eye blinks synchronize with musical beats during music listening

**DOI:** 10.1101/2025.02.10.637582

**Authors:** Yiyang Wu, Xiangbin Teng, Yi Du

**Affiliations:** State Key Laboratory of Cognitive Science and Mental Health, Institute of Psychology, Chinese Academy of Sciences, Beijing 100101, China; Department of Psychology, University of Chinese Academy of Sciences, Beijing 100049, China; Department of Psychology, The Chinese University of Hong Kong, Shatin, N.T., Hong Kong SAR 999077, China; Mind and Brain Institute, The Chinese University of Hong Kong, Shatin, N.T., Hong Kong SAR 999077, China; Chinese Institute for Brain Research, Beijing 102206, China

## Abstract

Auditory-motor synchronization, the alignment of body movements with rhythmic patterns in music, is a universal human behavior, yet its full scope remains incompletely understood. Through four experiments with 123 young non-musicians, integrating eye-tracking, neurophysiological recordings, white matter structural imaging, and behavioral analysis, we reveal a previously unrecognized form of synchronization: spontaneous eye blinks synchronize with musical beats. Blinks robustly synchronized with beats across a range of tempi and independently of melodic cues. EEG recordings revealed a dynamic correspondence between blink timing and neural beat tracking. Individual differences in blink synchronization were linked to white matter microstructure variation in the left arcuate fasciculus, a key sensorimotor pathway. Additionally, the strength of blink synchronization reflected the modulation of dynamic auditory attention. These findings establish blink synchronization as a novel behavioral paradigm, expanding the auditory-motor synchronization repertoire and highlighting the intricate interplay between music rhythms and oculomotor activity. This discovery underscores a cross-modal active sensing mechanism, offering new insights into embodied music perception, rhythm processing, and their potential clinical applications.

## Introduction

Auditory-motor synchronization—the coordination of body movements with auditory rhythms, such as tapping our feet or fingers to the musical beat—represents a universal human behaviour. This phenomenon, however, is relatively rare among species closely related to humans, suggesting the evolution of specialized auditory-motor neural circuitry within the human brain[1, 2]. Given that deeper understanding of behaviours often heralds deeper insights into neural systems underlying them[3], studying auditory-motor synchronization not only advances our knowledge of its distinct neural circuitry but may also offer valuable perspectives on brain disorders. Rhythm and timing deficits, for instance, are key vulnerabilities in neurodevelopmental conditions like autism spectrum disorder and developmental language disorder[4]. Moreover, auditory-motor synchronization has already proven effective in therapeutic and educational contexts for addressing motor or language disorders[5, 6]. Despite extensive research, the full spectrum of auditory-motor synchronizing behaviours remains incompletely characterized. In this study, we unveil a previously unrecognized form of synchronization: spontaneous eye blinks that synchronize with musical beats. This discovery highlights a novel functional and neural linkage between cortical auditory processing and oculomotor mechanisms.

Auditory-motor synchronization is typically associated with overt actions such as finger tapping, clapping, dancing, or whispering, where body movements align with rhythmic auditory stimuli[7, 8]. These voluntary movements are thought to engage both cortical and subcortical pathways, involving interactions between the auditory cortex, motor and premotor cortex, basal ganglia, and cerebellum[9–13]. According to the active sensing[14, 15] and predictive coding[16, 17] models in auditory contexts, and the “action simulation for auditory prediction” (ASAP) hypothesis[18] for musical beat perception in particular, motor recruitment helps refine temporal predictions of auditory patterns, optimize attention allocation, and improve auditory perception[19–21].

However, synchronized behaviours may extend beyond overt voluntary movements to include more subtle, involuntary motor actions. Music has profound effects on emotions[22], often engaging the dopamine system[23], modulating cognitive processes[24], and influencing peripheral autonomic systems[25]. These effects raise an intriguing question: could music rhythmically entrain spontaneous behaviours such as eye blinks—small but frequent motor actions with established links to dopaminergic function[26], attention[27] and cognitive effort[28]. While prior research has demonstrated that pupil dynamics can correlate with rhythmic structures[29], and that saccadic eye movements can synchronize with auditory rhythms[30], the relationship between eye blinks and musical rhythms remains unexplored. Eye blinks have been studied in relation to emotion[31], attention[32], and subjective states[33] during music listening, but their potential synchronization with musical rhythms presents a novel avenue for investigation.

In this study, we investigate the synchronization of eye blinks with musical beats, a fundamental musical rhythm. Through a combination of behavioural, neurophysiological and neuroimaging analyses, we provide robust evidence for this phenomenon, uncover its neural and structural substrates, and explore its functional role. Participants listened to Western classical music with highly regular beat patterns while their eye blinks were monitored, without any specific instructions regarding blinking. Electroencephalogram (EEG) responses were recorded to examine how musical structures are encoded in the brain and their correlation with blink activity. Furthermore, we analyzed the microstructural properties of white matter tracts connecting the frontal, parietal and auditory regions to identify structural differences linked to individual variation in auditory-oculomotor synchronization. Finally, we investigated the functional significance of this phenomenon, hypothesizing that stronger blink synchronization would correlate with better performance for events that aligned with the anticipated beat, consistent with the dynamic attending theory (DAT) [34–36]. Overall, this research reveals a new form of auditory-motor synchronization—blink synchronization—in music listening, and offers a mechanistic explanation through integrated behavioural, neural, and structural evidence. This work broadens our understanding of the intricate relationship between auditory rhythms and oculomotor behaviour, while also laying the groundwork for future studies on its potential clinical applications, especially as an objective and implicit biomarker for diagnosing dopamine-related and neurodevelopmental disorders.

## Results

### Eye blinks spontaneously synchronize with musical beats

In Experiment 1, 30 non-musicians listened to 10 Bach chorales at 85 beats per minute in both the original and reverse versions, while EEG and eye-tracking recordings simultaneously. In the reverse version, the sequence of beats was inverted from end to beginning while maintaining temporal regularity at the beat level and the original’s acoustic properties (Fig 1B and 1C). This design ensured the reverse version remained an unfamiliar control. Each musical piece was repeated three times consecutively to collect sufficient data on a single piece and measure the effect of repetition. The experimental procedure is summarized in Fig 1A and 1B. The acoustic modulation spectra of all musical pieces are shown in Fig 1C, demonstrating prominent frequency components of musical beat structures (beat rate: 1.416 Hz). To ensure that the reversal manipulation of harmonic progressions did not affect participants’ liking of the music, we compared the liking ratings for the original and reverse versions. No significant difference in liking ratings was found between the two versions (*t*_(29)_ = 0.715, *p* = 0.480, Cohen’s *d*□ = 0.131; Fig 1D), suggesting that preferences were unlikely to account for any differences in blink or neural signals observed later.

**Fig 1.**
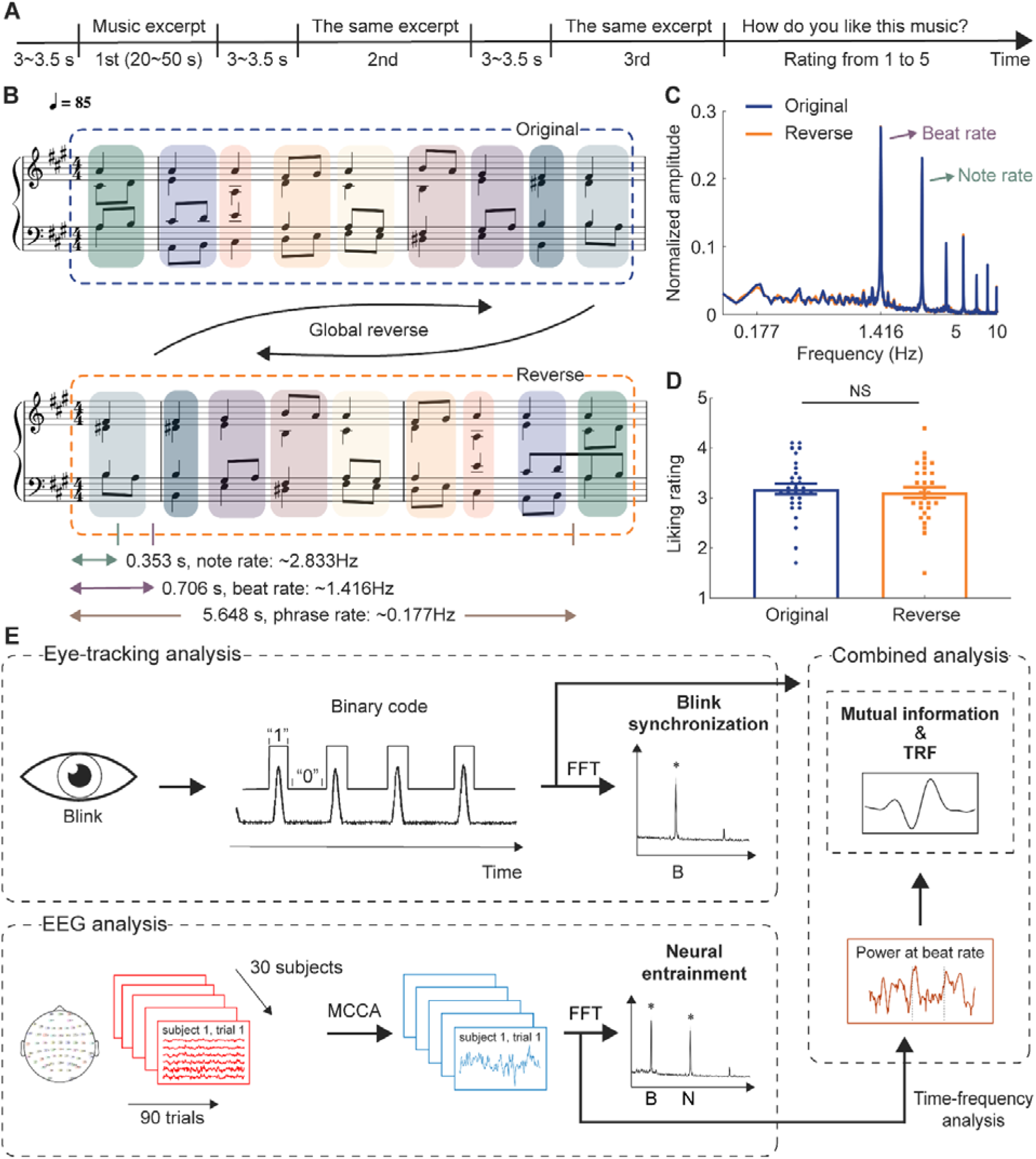
Design and data analysis pipelines of Experiment 1. **(A)** Each music excerpt was presented three times consecutively, during which EEG and eye-tracking data were recorded. After each trial, participants rated their liking of the excerpt. **(B)** Example of music excerpts. The tempo is 85 beats per minute: each note is ∼0.353 s (2.833 Hz), each beat is ∼0.706 s (1.416 Hz), and each phrase is ∼5.648 s (0.177 Hz). The reverse version (bottom) was created by reversing the beat order of the original excerpts (top). **(C)** Modulation spectra of the stimuli showed peaks at the note and beat rates as well as their harmonics for both versions. **(D)** Liking rate showed no significant difference between the two versions. **(E)** Overview of the analysis. Blinks were recorded via an eye tracker and converted into binary time series. The synchronization value was measured by the fast Fourier transform (FFT) of the blink signals. EEG was recorded with 64 electrodes, and multiway canonical correlation analysis (MCCA) extracted the most common components across subjects. Denoised signals were used to measure neural entrainment to beats and notes. Neural signal power at the beat rate was obtained using a wavelet transform. Mutual information and temporal response function (TRF) measured the relationship between blink signals and neural power at the beat rate.

Next, we measured blink synchronization with musical beats and neural entrainment to beats (Fig 1E). We quantified continuous eye-tracking data by converting it into binary time series to identify the timing of eye blinks. Fast Fourier transform was then applied to the blink signals, and amplitude spectra of eye blinking were derived. Similarly, we extracted music-related EEG components (see Methods for details) and derived amplitude spectra to examine whether neural components corresponded to musical beats, a routine analysis of neural entrainment to musical rhythms.

To our surprise, while neural entrainment to musical beats has been well documented[37–42], a similar finding was shown in the spectra of eye blinking: eye blinks spontaneously synchronized with musical beats, even without any instructions to do so. The blink synchronization with musical beats was significant, as demonstrated by the spectral peaks of eye blinking dynamics exceeding the thresholds (*p* < 0.01) created through a surrogate test on the group-averaged data (Fig 2A). We further derived the corrected amplitude within the significant frequency range for further analyses, which spanned from 1.416 Hz to 1.433 Hz.

**Fig 2.**
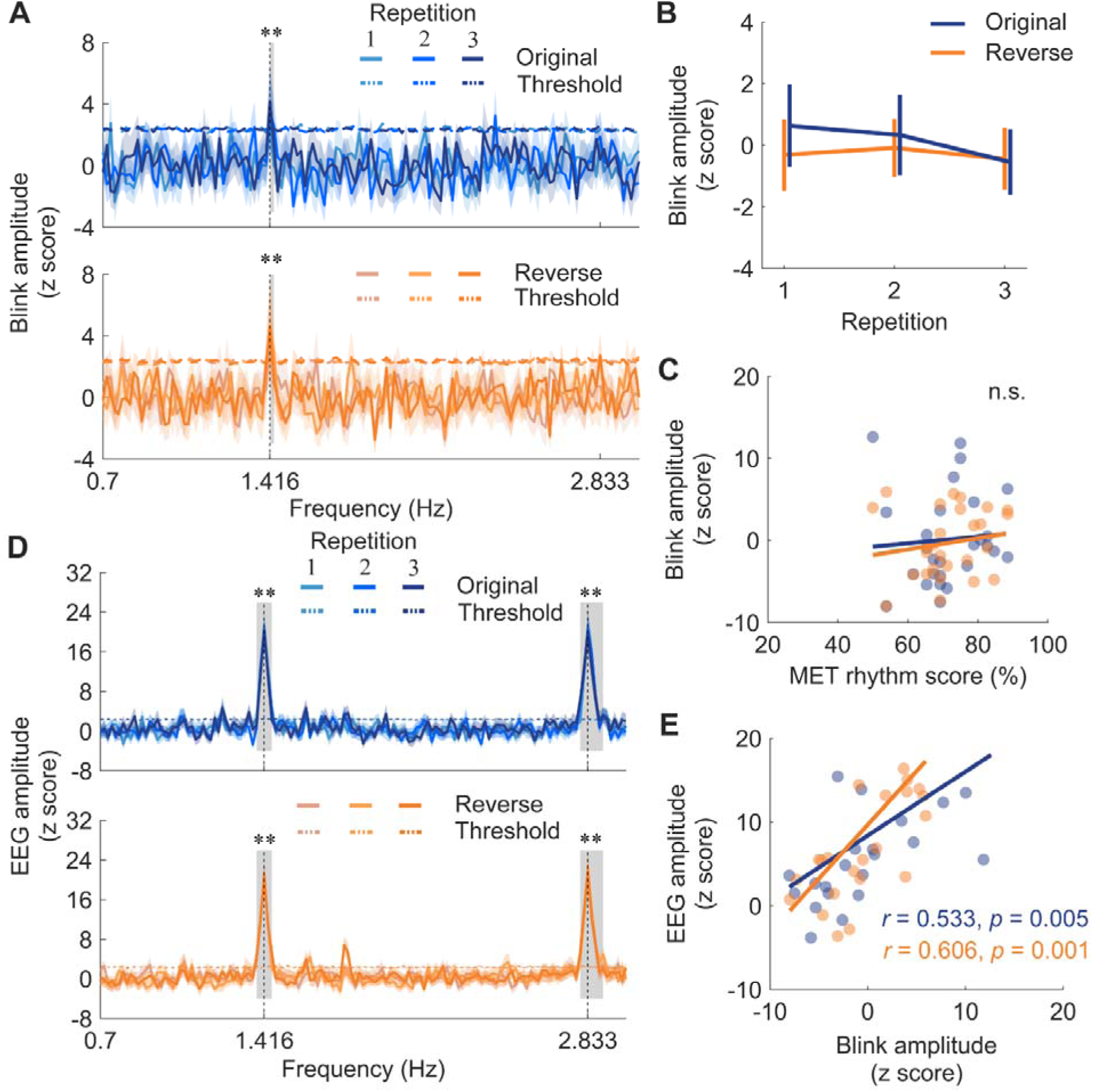
Blink and neural synchronization to musical beats. **(A)** Blink amplitude spectra for the two versions and repetitions, with horizontal dashed lines showing the threshold from the surrogate test (*p* < 0.01). Shaded areas represent ± one SEM across participants (n = 30). Gray boxes indicate the frequency ranges where amplitude was above the threshold. The blink amplitude showed a salient peak around the beat rate. **(B)** Blink amplitude within significant frequency ranges around the beat rate, with error bars denoting ± one SEM. Neither version nor repetition had a significant effect on blink amplitude. **(C)** Correlation between the MET rhythm score and blink amplitude, showing no significant relation (n.s., not significant). Colored dots represent individuals. **(D)** Neural amplitude spectra for the two versions, with shaded areas representing ± one SEM (n = 30). Horizontal dashed lines represent surrogate test thresholds (*p* < 0.01). Gray boxes indicate the significant frequency ranges. Salient peaks appeared at the note and beat rates for both versions. **(E)** Correlation between blink amplitude and neural amplitude at the beat rate. Blink amplitude was positively correlated with neural amplitude for both versions. Colored dots represent individuals. ** *p* < 0.01.

This finding, though surprising and perhaps entirely new, was robustly observed across both versions of musical pieces and all three repetitions. This is further echoed by a histogram of raw intervals between two consecutive blinks (S1 Fig), which shows a clear peak of intervals between one and two beats.

To assess the effects of different versions of musical pieces and repetition on blink synchronization, we conducted a two-way repeated-measures ANOVA, with version (original or reverse) and repetition (1, 2 or 3) as main factors, on the corrected amplitude within the significant frequency range (Fig 2B). We did not find any significant main effects nor interaction (version: *F*_(1,26)_ = 0.398, *p* = 0.534, *η_p_^2^* = 0.015; repetition: *F*_(2,52)_ = 0.419, *p* = 0.660, *η_p_^2^* = 0.016; interaction: *F*_(2,52)_ = 0.238, *p* = 0.789, *η_p_^2^* = 0.009). This lack of significance may stem from the fact that the beat structure was well preserved in different versions and was easy for participants to follow in each condition. As a result, even though blink synchronization to the beats was evident, no significant differences were found between repetitions or between the original and reverse versions.

We then tested whether blink synchronization performance correlates with musical ability measured by the rhythm subtest of the Musical Ear Test (MET)[43]. The corrected amplitude within the significant frequency range were averaged across repetitions for each version and correlated with the MET rhythm score to examine the effect of musical rhythmic ability. No significant correlations were found for either version (original: *r*_(27)_ = 0.072, *p* = 0.720; reverse: *r*_(27)_ = 0.168, *p* = 0.403; Fig 2C). This lack of correlation is likely due to the fact that the beat structure in our stimuli was relatively easy to synchronize to, as noted earlier.

### Neural responses entrained to musical structures

In Experiment 1, we also replicated the established phenomenon of neural entrainment to musical beats. In line with previous studies[37–42], robust neural entrainment to beats was evident (Fig 2D). After preprocessing raw EEG data and removing independent components associated with eye blinks, eye movements, and heartbeat using an independent component analysis (ICA) algorithm, we applied multiway canonical correlation analysis (MCCA) to extract components linked to music listening (see Methods). This approach enabled the extraction of auditory neural signals specific to music listening and facilitated the analysis of single-trial data for each musical piece and repetition. We then derived amplitude spectra of neural signals corresponding to music listening, and conducted the surrogate test to extract the corrected amplitude within the significant frequency ranges (at beat rate, 1.383 – 1.45 Hz; at note rate, 2.8 – 2.9 Hz) for both beat and note rates for further analyses.

To examine the effects of version and repetition, we performed a two-way repeated-measures ANOVA for beat and note rates separately (S2 Fig). At the note rate, the analysis showed no main effect of version (*F*_(1,26)_ = 0.948, *p* = 0.339, *η_p_^2^* = 0.035) nor interaction (*F*_(2,52)_ = 0.718, *p* = 0.493, *η_p_^2^* = 0.027), but a main effect of repetition (*F*_(2,52)_ = 4.078, *p* = 0.023, *η_p_^2^* = 0.136). At the beat rate, neural tracking of beats was unaffected by either factor or their interaction (version: *F*_(1,27)_ = 0.196, *p* = 0.661, *η_p_^2^* = 0.007; repetition: *F*_(2,54)_ = 1.566, *p* = 0.218, *η_p_^2^* = 0.055; interaction: *F*_(2,54)_ = 1.334, *p* = 0.272, *η_p_^2^* = 0.047). Additionally, neural entrainment to notes and beats correlated with the MET rhythm score solely for the reverse version (note rate: *r*_(27)_ = 0.445, *p* = 0.020; beat rate: *r*_(28)_ = 0.454, *p* = 0.015). This suggests that higher musical ability is necessary for effectively tracking beat structures in the reverse version, possibly due to the unfamiliar harmonic progressions. Consequently, this version demands enhanced rhythmic skills to accurately follow the music’s beat patterns upon first exposure.

### Neural entrainment to musical beats corresponds to eye blinks

Subsequently, we examined the correspondence between blink synchronization and neural entrainment to beats. It is shown that the strength of neural entrainment correlates with behavioural measures of sensorimotor synchronization skills[44]. Extending this, the current study discovered that blink synchronization performance was significantly correlated with neural entrainment (original: *r*_(26)_ = 0.533, *p* = 0.005; reverse: *r*_(26)_ = 0.606, *p* = 0.001; Fig 2E). Given the distinct scalp topographies of the extracted neural components and eye-related components (S3 Fig), the blink-neural relation observed here cannot simply be attributed to blink artifacts.

To further investigate the underlying mechanism, we examined their relationship through mutual information (MI) and eye-blink triggered temporal response function (TRF). MI between eye blink timing series and neural signals across entire musical pieces indicates whether the brain encodes information from the blinks. Given the blink-neural relationship was specific to the beat rate, we conducted a wavelet transform on EEG signals to derive the power of neural responses at the beat frequency, and then calculated the MI between the EEG power and blink signals. Indeed, compared to surrogate data, empirical MI values were significantly larger for all repetitions in the original version (*ps* < 0.01; Fig 3A). However, the reverse version yielded a significant result only in the third presentation (*p* = 0.483, *p* = 0.092 and *p* = 0.005 for the first, second and third presentations, respectively). A two-way repeated-measures ANOVA revealed a significant main effect of version (*F*_(1,27)_ = 4.396, *p* = 0.046, *η_p_^2^* = 0.140; Fig 3B), showing higher MI in the original version. It also revealed a significant main effect of repetition (*F*_(1.629,43.979)_ = 9.767, *p* = 0.001, *η_p_^2^* = 0.266), with higher MI on the second and third presentations than the first (*p* = 0.002 and *p* = 0.010, second and third presentations respectively). The interaction was not significant (*F*_(2,54)_ = 1.479, *p* = 0.237, *η_p_^2^* = 0.052).

**Figure 3.**
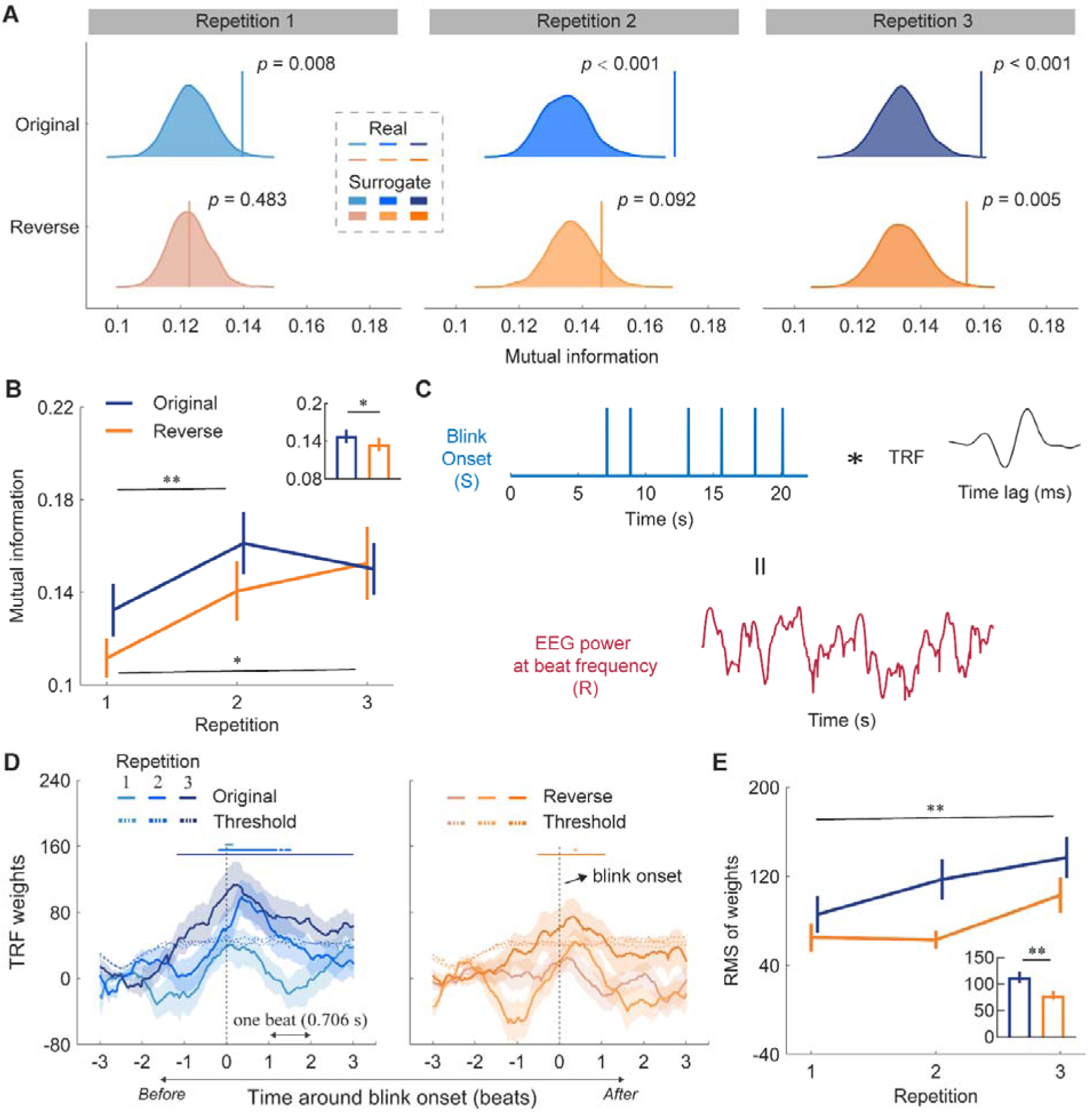
Correspondence between blinks and neural signals. **(A)** Mutual information (MI) between blink signals and beat-rate EEG power. Colored distributions represent surrogate MI values derived from circularly shifted EEG data. Colored vertical lines indicate the observed MI values. **(B)** Averaged MI for each version and repetition. The bar plot shows the mean MI values for both versions. Error bars denote ± one SEM. MI increased with repetition and was higher in the original version. **(C)** Temporal response function (TRF) analysis of EEG and eye blinks. Blink onsets were extracted from the blink signals and regressed against the EEG signals, resulting in the blink onset TRF. **(D)** TRF of beat-rate EEG power using blink onset as the regressor. Shaded areas represent ± one SEM across participants (n = 30). Vertical dashed lines mark blink onset. Horizontal axis covers three beats before and after blink onset. Horizontal dashed lines show permutation test thresholds (*p* < 0.01). Horizontal solid lines at the top indicate significance. Notably, TRF weights increased significantly before blink onset. **(E)** Root mean square (RMS) of TRF weights at the peak. The bar plot shows the mean TRF weights for both versions. Error bars denote ± one SEM. RMS of weights was larger in the original version. * *p* < 0.05, ** *p* < 0.01.

Interestingly, MI results suggest that repetitions or listeners’ familiarity with musical pieces modulated neural coupling to eye blinks, despite no such effect being observed on either neural entrainment (S2 Fig) or blink synchronization (Fig 2B), individually. This dependency was evident across all repetitions of the original version but only emerged during the third repetition of the reverse version. This discrepancy between the two versions, despite featuring identical beat structures, implies that effective anticipation, driven by normal harmonic progressions in the original version, helps bind blink and neural responses. Furthermore, the repetition effect underscores that blink synchronization transcends mere reflexive responses to sensory stimulation.

### Neural entrainment to musical beats is time-locked to blink onset

Next, we investigated how the brain encodes eye blinks over time. Employing TRF methodology, we extracted blink onset as the specific feature and examined the fluctuations in neural power at the beat rate before and after eye blinks, as illustrated in Fig 3C. Remarkably, neural responses were time-locked to blink onset, with an increase in neural responses, as reflected in TRF weights, prior to blink onset (Fig 3D). Specifically, for the original version, TRF weights were significantly higher than the surrogate-derived threshold before blink onset during the second and third presentations. Similarly, for the reverse version, TRF weights exceeded the threshold before blink onset during the third presentation. The repetition effect observed here echoes the MI results. A two-way repeated-measures ANOVA on the root mean square (RMS) of TRF weights at the response peak found a significant main effect of version (*F*_(1,27)_ = 10.420, *p* = 0.003, *η_p_^2^* = 0.278; Fig 3E), suggesting higher RMS of weights for the original version. It also showed a significant main effect of repetition (*F*_(2,54)_ = 4.799, *p* = 0.012, *η_p_^2^* = 0.151), with higher weights during the third presentation than the first presentation (*p* = 0.004, Bonferroni corrected), indicating a gradual formation of prediction for blink onset in the brain.

Thus far, we have established that blink synchronization—wherein eye blinks track musical beats—robustly corresponds with neural entrainment to beats. Moreover, blink synchronization transcends mere passive reflex to sensory stimuli, and the blink-neural relation is modulated by cognitive factors such as repetition and reversal manipulations.

### Microstructure of the left arcuate fasciculus is associated with blink synchronization

Previous studies have revealed notable individual differences in auditory-motor synchronization abilities, which are linked to variations in white matter microstructural properties of sensorimotor pathways[45–47]. In the current study, we assessed the relationship between white matter microstructural properties and blink synchronization performance to further explore the structural variability associated with blink synchronization. Given the established positive correlation between fractional anisotropy (FA) of the arcuate fasciculus (AF) and auditory-motor synchronization[45–47], we hypothesized that individuals showing better blink synchronization performance would display stronger structural connectivity in this key sensorimotor pathway.

Firstly, we excluded 3 outliers and averaged the blink response amplitudes across two versions to obtain a global measure of blink synchronization. We then performed a median split based on the mean blink amplitude to divide participants into two groups: 13 high synchronizers and 13 low synchronizers (excluding the median, Fig 4A). The blink amplitude spectra for the two groups showed a significant difference around the beat frequency (S4 Fig). However, no significant difference was observed between the two groups concerning their MET rhythm scores (*t*_(24)_ = -1.162, *p* = 0.257, Cohen’s *d*┚ = -0.458; Fig 4B). Notably, participants categorized as high synchronizers based on blink amplitude also showed stronger neural entrainment at the beat rate compared to low synchronizers (*u* = 141, *p* = 0.003; Fig 4C).

**Figure 4.**
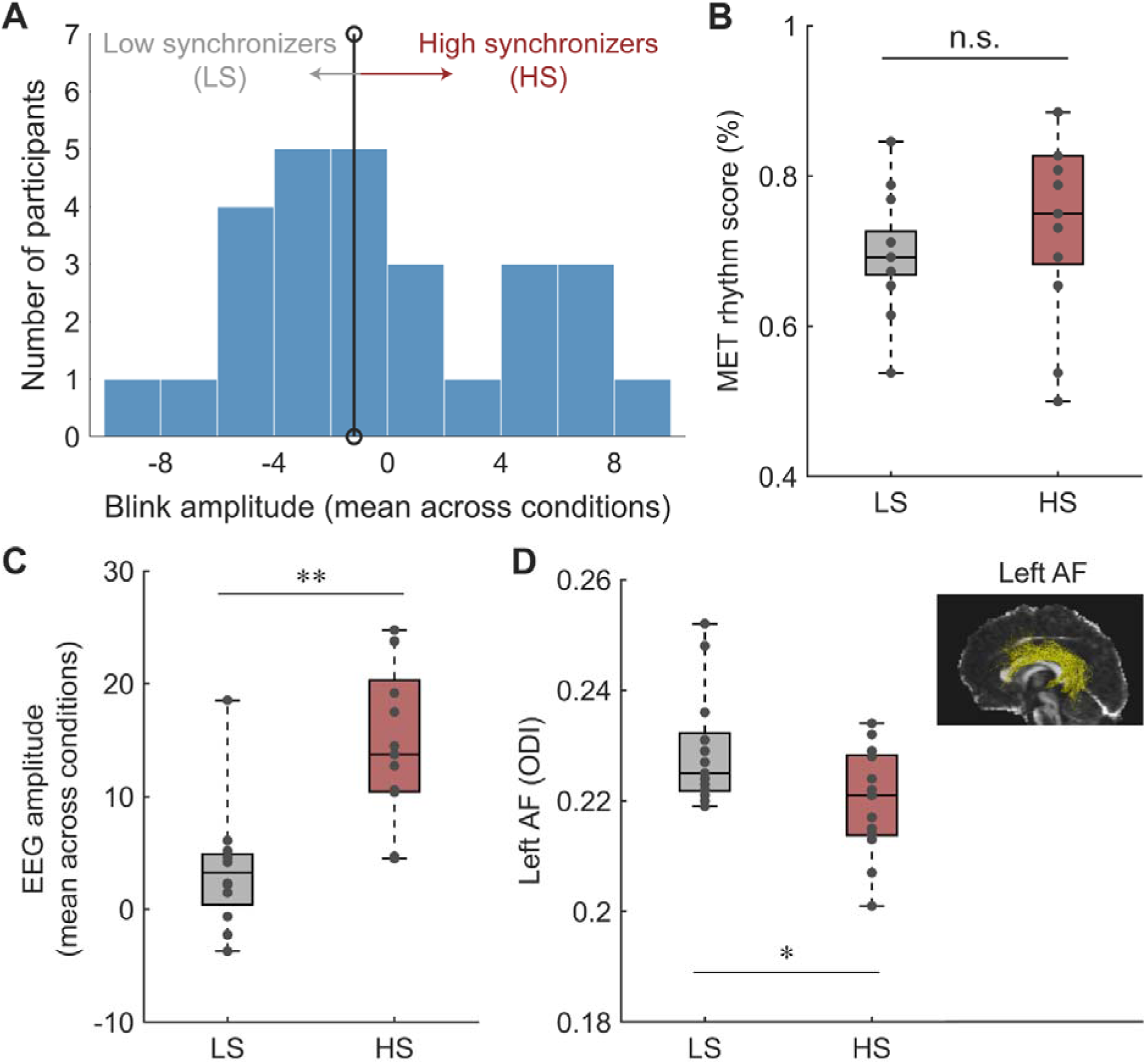
Individual differences in blink synchronization and microstructural property. **(A)** Histogram of blink amplitude averaged across conditions. Participants were divided into two groups based on a median split of their blink amplitude: low synchronizers (below the median, n =13) and high synchronizers (above the median, n =13). The vertical line marks the median. **(B)** Boxplots of MET rhythm scores for the two groups. No significant difference in musical ability was found between the groups. **(C)** Boxplots of neural entrainment to beats (EEG amplitude averaged across conditions) for the two groups. Neural entrainment to beats was stronger in high synchronizers than in low synchronizers. **(D)** Boxplots of the orientation dispersion index (ODI) of the left arcuate fasciculus (AF). The ODI was higher in low synchronizers than in high synchronizers. The inserted plot displays the tractography of the left AF in one typical participant. * *p* < 0.05, ** *p* < 0.01, n.s., not significant.

We then acquired diffusion-weighted imaging (DWI) data to quantify potential differences in white matter tracts connecting frontal, parietal and auditory regions that might distinguish the two groups in terms of blink synchronization. In addition to the AF, we also focused on the superior longitudinal fasciculus (SLF), that links auditory and dorsal premotor regions through parietal regions (see Fiber tractography in Methods) and have been implicated in auditory-motor synchronization and beat perception[18]. We extracted three microstructural indices—FA from the diffusion tensor imaging (DTI) model, neurite density index (NDI) and orientation dispersion index (ODI) from the neurite orientation dispersion and density imaging (NODDI) model—as well as their lateralization indices (LI) for bilateral SLF and AF segments to explore the relationship between blink synchronization and white matter properties. Independent t-tests on each microstructural index and its corresponding LI revealed a significant group difference only in the ODI of the left AF, with low synchronizers showing higher ODI than high synchronizers (*t_(24)_* = 2.272, *p* = 0.032, Cohen’s *d*□= 0.910, Fig 4D). This suggest that poorer blink synchronization may be linked to more dispersed, less efficient structural connectivity in the left AF, impairing beat synchronization, while better performance may be associated with more organized and aligned white matter fibers in the left AF. No significant group differences were observed for other microstructural indices (see S1 Table for full statistics).

### Eye blinks are primarily entrained by temporal patterns in music

Despite our confirmation of this novel behaviour, several questions remain, particularly regarding which musical elements drive this behaviour. Specifically, does blink synchronization depend on pitch information or melodic patterns, or is it solely driven by temporal patterns in tone sequences deprived of melodic cues? Additionally, given that rhythmic rate has been shown to influence people’s abilities to synchronize with auditory-motor activities, such as tapping, whispering or clapping in synchrony with auditory stimuli[48, 49], does musical tempo also modulate blink synchronization?

To address these questions, in Experiment 2, we compared blink synchronization to both music and static tone sequences across three tempi (66, 85, and 120 beats per minute) (Fig 5A). We generated ten new tone sequences maintaining the temporal structures of the original musical pieces but lacking pitch information or melodic cues. Similar to Experiment 1, 30 non-musicians listened attentively to each auditory stimulus and pressed the spacebar as soon as possible upon detecting a timbre deviant, while their eye movements were recorded. Trials containing timbre deviants were excluded from analyses.

**Figure 5.**
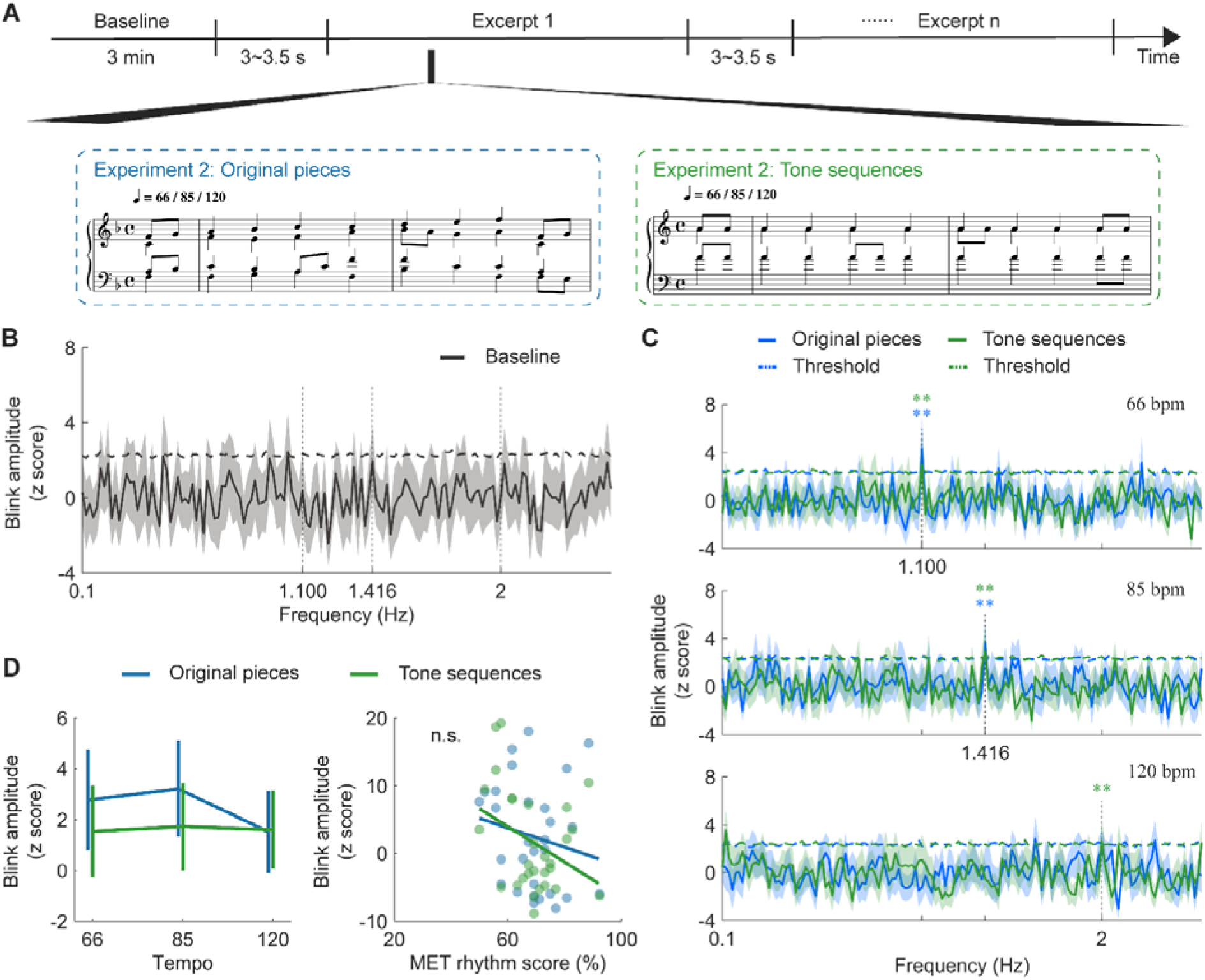
Blink synchronization in response to rhythmic sequences. **(A)** Experimental paradigm for Experiment 2: participants listened to both original musical pieces and tone sequences while their eye movements were recorded. **(B)** Blink amplitude spectra for the baseline (no music), with horizontal dashed lines representing the surrogate test threshold (*p* < 0.01). Shaded areas represent ± one SEM across participants (n = 30). **(C)** Blink amplitude spectra for two stimuli types at three tempi, with horizontal dashed lines showing the surrogate test threshold (*p* < 0.01). Vertical dashed lines indicate the frequency points corresponding to the beat rates. Shaded areas represent ± one SEM. Blink amplitude showed salient peaks at the beat rate across stimuli types and tempi, except at 120 beats per minute for original pieces. ** *p* < 0.01. **(D)** Left: Blink amplitude at the beat rate. Error bars denote ± one SEM. Stimuli type and tempo had no significant effect on blink responses. Right: No significant correlation was found between MET rhythm score and blink amplitude. Colored dots indicate individuals (n.s., not significant).

A key difference from Experiment 1 was the inclusion of a 3-minute rest period before the main experiment to determine whether eye blinks during idle state exhibit any temporal structures. A surrogate test on the baseline data yielded no significant peak (Fig 5B), confirming that the spectral peak observed in Experiment 1 was indeed music-related or at least stimulus-related.

In Fig 5C, we replicated the findings in Experiment 1, observing robust blink synchronization in Experiment 2. Importantly, musical tempo modulated blink synchronization, with spectral peaks aligning with the beat rates across stimulus types and tempi, except at the highest tempo (120 beats per minute) for original pieces. We extracted the corrected amplitude within the significant frequency ranges derived from the surrogate test for the three tempi: 66 beats per minute, 1.100 Hz; 85 beats per minute,1.416 Hz; and 120 beats per minute, 2 Hz. Then we tested the effects of stimuli type (original pieces or tone sequences) and tempo (66, 85 or 120 beats per minute) using a two-way repeated-measures ANOVA. Although blink synchronization appeared stronger for original pieces than tone sequences and decreased with increasing tempo, we found no significant main effects (stimuli type: *F*_(1,28)_ = 0.510, *p* = 0.481, *η_p_^2^* = 0.018; tempo: *F* = 0.352, *p* = 0.705, *η_p_^2^* = 0.012) or interaction effect (*F*_(2,56)_ = 0.182, *p* = 0.834, *η_p_^2^* = 0.006). Moreover, there was no significant correlation between blink synchronization and the MET rhythm score (original pieces: *r*_(29)_ = -0.287, *p* = 0.131; tone sequences: *r*_(29)_ = -0.337, *p* = 0.073; Fig 5D).

Taken together, the results in Experiments 1 and 2 suggest that blink synchronization is primarily entrained by the temporal patterns in music, rather than pitch changes or melodic cues. Eye blinks align with the timing of auditory events, supporting the idea that temporal structure is the key driver of synchronization.

### Blink synchronization facilitates pitch deviant detection

In Experiment 3, we investigated whether blink synchronization correlates with cognitive performance. While prior experiments have established a connection between blink synchronization and neural entrainment to beats, we further sought to determine if better blink synchronization corresponds to improved tracking of musical elements? To do so, we conducted a pitch deviant detection task where non-musicians (n = 31) listened to music segments selected from the original pieces. Each segment contained two phrases (16 beats in total), with a pitch deviant introduced in the second phrase. Participants were instructed to detect deviant tones as soon as possible, while their eye movements were recorded (Fig 6A).

**Figure 6.**
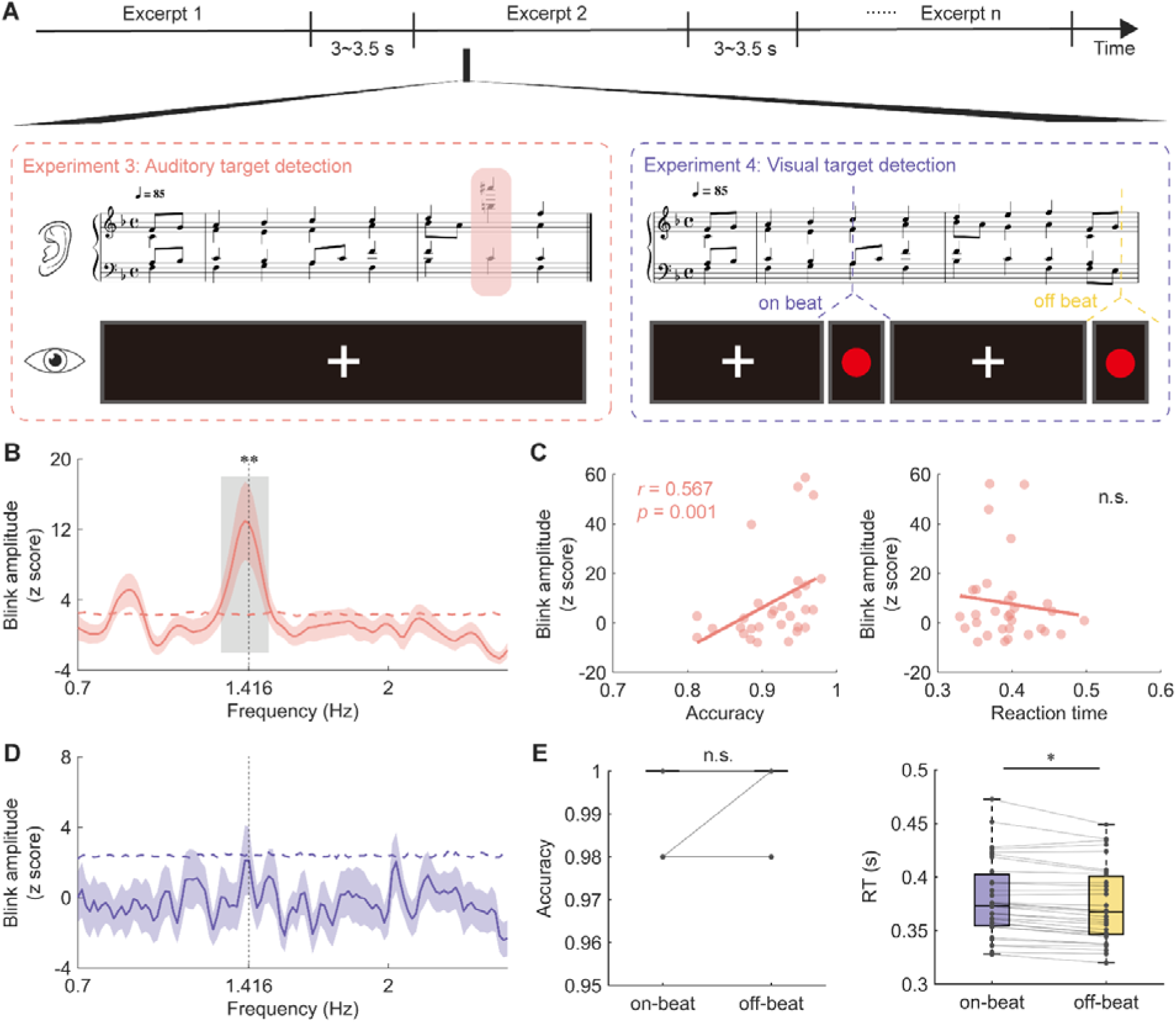
Functional significance of blink synchronization in behavioral performance. **(A)** Experimental paradigm for Experiments 3-4: participants performed a pitch deviant detection task (Experiment 3) or a visual target detection task (Experiment 4) while their eye movements were recorded. **(B)** Blink amplitude spectra in Experiment 3, with horizontal dashed line representing the surrogate test threshold (*p* < 0.01). Shaded area represents ± one SEM across participants (n = 31). The gray box indicates the frequency range above the threshold. The blink amplitude showed a salient peak around the beat rate. **(C)** Correlation between blink amplitude and behavioral indexes in Experiment 3. Blink amplitude was positively correlated with detection accuracy, but not with reaction time (n.s., not significant). Colored dots indicate individuals. **(D)** Blink amplitude spectra in Experiment 4 (n = 32), with no significant peak observed. **(E)** Group-averaged accuracy and reaction time in the on-beat (purple) and off-beat (yellow) conditions. Colored connecting lines represent individual participants. Participants responded significantly faster to off-beat visual target than to on-beat target. * *p* < 0.05, n.s., not significant.

We replicated the signature of blink synchronization to beats observed in earlier experiments (Fig 6B). We further found a positive correlation between blink synchronization strength and deviant detection accuracy (*r*_(30)_ = 0.567, *p* = 0.001; Fig 6C left). However, no significant association was observed between blink synchronization and reaction time (*r*_(31)_ = -0.072, *p* = 0.701; Fig 6C right). These results suggest that proficient blink synchronization may facilitate the detection of deviant pitch by aligning temporal attention with beat onset, implying a potential role of blink synchronization in a dynamic attending process.

### Blink synchronization disappeared when music is task-irrelevant

In Experiment 3, we suggested that blink synchronization reflects a dynamic attending process. Given that one of the functions of eye blinks is to regulate the flow of visual information, we next investigated blink synchronization in a cross-modal context in Experiment 4. The experimental procedure mirrored that of Experiment 3, except that 32 non-musician participants were required to detect a simple visual target—a red dot—presented on the screen while listening to music (Fig 6A). The deliberate simplicity of the visual task was designed to ensure participants’ attention primarily towards the music. We manipulated the temporal relationship between the visual target and musical beat onsets, resulting in two conditions: on-beat and off-beat. If blink synchronization reflects the modulation of visual sampling efficiency by musical rhythms, we would expect to observe a correlation between blink synchronization and behavioral performance as found in Experiment 3, and better performance, like higher accuracy or shorter reaction time, in the on-beat condition than the off-beat condition.

Despite these expectations, eye blinks did not even synchronize with musical beats in this task (Fig 6D). This observation is particularly noteworthy, as participants were not heavily engaged in the visual task, as evidenced by nearly 100% accuracy in both conditions (*t*_(27)_ = -0.812, *p* = 0.424, Cohen’s *d* □ = □ -0.153, Fig 6E left). A plausible explanation is that blink synchronization is a spontaneous behavior induced by active engagement in music listening, rather than a response to passive auditory stimuli. We further compared reaction time for detecting the visual target between on-beat and off-beat conditions using a two-sided paired t-test. Unexpectedly, responses were significantly slower in the on-beat condition (*t*_(30)_ = 2.417, *p* = 0.022, Cohen’s *d*□ = 0.434, Fig 6E right).

## Discussion

We’ve demonstrated through a series of experiments that oculomotor activity, specifically eye blinking, spontaneously aligns with regular musical beats during active music listening—a phenomenon we term blink synchronization, and explored its neural substrates and functional significance. As depicted in Fig 2, we discovered that eye blinks spontaneously synchronize with musical beats at 85 beats per minute. Moreover, we observed a correspondence between blink synchronization and neural entrainment to beats (Fig 3). On a structural level, we found that the microstructural connectivity of the left AF played an important role in the strength of blink synchronization (Fig 4). We then replicated this blink synchronization phenomenon across a broader range of tempi (66 – 120 beats per minute) and in auditory sequences without melodic cues (Fig 5), affirming its robustness. Furthermore, our investigation into the functional relevance of blink synchronization revealed intriguing findings. Experiment 3 revealed a correlation between stronger blink synchronization and improved performance on a pitch deviant detection task in a musical context. Experiment 4 using a visual detection task highlighted the impact of task relevance on blink synchronization to music (Fig 6).

The spontaneous blink rate has long been recognized as a vital indicator of cognitive and neural functions[50]. Recent research has begun to explore the link between eye blinks and music listening, aiming to uncover the cognitive complexities of human music processing related to emotion[31], attention[32], and subjective states[33]. However, a direct connection between blinks and beat perception remains elusive. While motor behaviours such as finger tapping or dancing are commonly studied to understand musical rhythm and beat perception, an intriguing question remained: can music implicitly entrain blink activity? Our discovery of blink synchronization established a direct link between eye blinks and musical beat perception in a naturalistic listening setting, introducing a novel type of spontaneous music-synchronizing behaviour. This is especially interesting because when people tap their feet or nod their heads in sync with the music, they are typically aware of it. In contrast, although we did not assess participants’ self-awareness of their blinking behaviour, none reported consciously blinking to the beat after the experiment. The fact that autonomic oculomotor activity can track musical beat may reflect the evolution-rooted primitive instinct in music rhythm processing[18]. In other auditory contexts, eye blinks have been shown to track high-level linguistic and non-linguistic structures, reflecting a modulation of temporal attention[51]. Future works should investigate whether blink synchronization extends to higher-level musical structures, such as meter or phrase, to clarity the cognitive processes and mechanisms involved and differentiate oculomotor synchronization in music from that in speech processing.

Our results also provide compelling evidence for brain-eye coordination in beat perception, with blink activity and neural entrainment activity coupled during music listening (Fig 3A and 3B). Specially, eye blinks were represented in cortical responses, with a time-locked neural response gradually preceding blink onset (Fig 3D and 3E), suggesting a neural prediction of upcoming blinks refined through exposure. Aligning with models such as predictive coding[16, 17], active sensing[14, 15], and the ASAP hypothesis[18], our findings suggest that motor cortical oscillations not only synchronizes with incoming musical beats, but also actively predicts impending blink activities. By comparing the expected and actual timing of blinks, the brain continuously updates its predictions, leading to improved synchronization between blinks and neural signals. This enhanced synchronization may, in turn, facilitate more efficient auditory beat processing.

Moreover, we found that the microstructural integrity (indexed by ODI) of the left AF, a tract connecting the auditory cortex and frontal motor areas, served as the structural basis for individual differences in blink synchronization. Lower ODI, which indicates less fiber dispersion and higher coherence, was associated with better blink synchronization performance (Fig 4D). ODI derived from the NODDI model quantifies the angular variation of neurites and provides a more precise measure of neurite dispersion than FA[52]. A series of studies have reported a decrease in FA with an increase in ODI of white matter, and the changes in neurite geometry were associated to cognitive impairment[53–55]. Our result aligns with previous findings showing that larger FA value of the AF supports higher auditory-motor synchrony[45–47]. Our findings underscore the crucial role of microstructural differences in the sensorimotor pathway connecting auditory and frontal motor regions in influencing individual variations in auditory-motor synchronization across auditory stimuli and motor effectors.

Notably, we did not find significant blink synchronization at a tempo of 120 beats per minute for the musical stimuli (Fig 5C). This implies a rate limit for blink synchronization, a phenomenon also observed in finger tapping and covert synchronization studies[49, 56]. Given differentially preferred tempo of rhythmic behaviours among motor effectors[48], further research is required to determine the optimal tempo range for robust blink synchronization to musical beats. Interestingly, blink synchronization was restored when original musical pieces were replaced with tone sequences at 120 beats per minute. This result is consistent with a recent study showing that repeated acoustic units facilitate auditory-motor synchronization in a subgroup of the population[57], inspiring us to take acoustic features into consideration in future experiments.

In Experiment 3, we found that blink synchronization performance was positively correlated with the detection accuracy of pitch deviant that appeared at the anticipated beat (Fig 6C), consistent with previous findings[19, 20, 58]. This result supports the DAT[34–36] and the active sensing model[14, 15], suggesting that, similar to other motor behaviours, synchronized blinks help entrain attention to musical beats, enhancing the precision of temporal prediction, thereby facilitating the detection of pitch deviants. However, in Experiment 4, blink synchronization disappeared when the music was task-irrelevant (Fig 6D), suggesting that auditory rhythms only entrain blinks when they are task-relevant and well-attended. A prior study has demonstrated that gaze aligns with speech dynamics and the strength is modulated by selective attention[59]. In our task, participants were instructed to focus on visual stimuli, which may have inhibited the processing of auditory information, resulting in the absence of blink synchronization. Additionally, the ceiling effect in task performance may have obscured active sensing. Further experiments are needed to explore the effect of synchronized blinks on cross-modal temporal modulation using more challenging tasks.

The delayed response to on-beat visual target (Fig 6E) was notable and seemingly contradictory to the DAT[34–36], which suggests that periodic auditory rhythms entrain attention to specific time points to optimize perception, yielding faster reaction times for stimuli presented in synchrony with the rhythm. Our results instead suggest a cross-modal interaction in dynamic attending. When visual targets appeared on the beat, participants may have focused more on the auditory beat event, delaying visual responses. In contrast, the off-beat condition likely allowed for more flexible attention allocation, leading to faster responses. Moreover, while most studies show that unattended musical rhythms facilitate perception and memory[60–62], other find that unattended musical rhythms either impair or have no effect on perception[63–65]. This disparity is likely due to the strength difference of temporal expectations provided by musical rhythms. In our study, the intermixed design in which on-beat and off-beat targets occurred within the same block, likely diminished the rhythmic effect on target detection. Our result may also reflect a post-blink boost effect, where blinks help reformat visual information and enhance perceptual sensitivity after blinks, as demonstrated in previous research[66, 67]. As illustrated in S5 Fig, blink rates were more likely to peak around the beat onset, potentially facilitating the perceptual processing of off-beat targets.

Musical beat perception likely engages a subcortical-cortical interaction between the basal ganglia and premotor areas[68]. While some studies have linked the premotor areas to peripheral oculomotor activities in non-human animals[69, 70], other studies on eyeblink conditioning have identified a close relationship between the basal ganglia and eye blinks[71, 72]. It is plausible that the basal ganglia, stimulated by musical beats, signals the frontal eye field and oculomotor muscles, prompting eye blinks via a subcortical-cortical-peripheral pathway. This echoes the finding shown in Fig 3D, where neural entrainment to beats predicted eye blinking activities. Nevertheless, the neural circuit underlying this phenomenon needs future investigations.

The discovery of blink synchronization marks a significant stride in our comprehension of auditory-motor synchronization, yet this study has certain limitations that present avenues for future exploration. First, the participant sample, consisting primarily of young non-musicians with normal hearing, should be expanded to broader populations. Including musicians can reveal how musical training influences blink synchronization, while diverse age groups could help assess developmental trajectories. Given that impaired rhythm processing is a risk factor for neurodevelopmental disorders[4], blink synchronization during music listening may have great potential to serve as an implicit and feasible diagnostic and interventional biomarkers for children, compared to explicit paradigms like finger tapping. Second, the exclusive use of Bach’s chorales, noted for high temporal regularity and representation of Western classical music, suggests a limitation in musical diversity. Investigating blink synchronization in response to music featuring irregular rhythms, syncopation, and a variety of musical genres and cultural backgrounds is crucial for understanding the universality of this phenomenon and its dependency on rhythmic structures. Furthermore, given that pupil dynamics[29] and saccades[30] are found to implicated in processing auditory rhythms and musical structures, the coordination between blinks, pupil size and saccades in beat perception warrants further investigation to advance our understanding of eye movements in music listening. Finally, future neuroimaging research should examine the neural underpinnings of blink synchronization, particularly focusing on the basal ganglia, frontal eye field and premotor areas.

In conclusion, our study sheds light on the relationship between music listening and oculomotor activity, specifically the synchronization of eye blinks with musical rhythms. Our findings establish blink synchronization as a novel, distinctive, spontaneous auditory-motor synchronization behaviour during music listening, and reveal its neurophysiological and structural correlates. By linking active sensing and ASAP frameworks to blink synchronization, we gain deeper insights into the coordination of auditory, motor, and visual systems, thereby enriching our knowledge of cross-modal interaction and embodied musical perception.

## Materials and methods

### Participants

A total of 123 young adults (70 women; mean age, 22.67 ± 2.98 years, ranging from 18 to 34 years old) with normal hearing (thresholds ≤ 20 dB SPL from 125 to 8000 ≤ Hz) took part in the study. Thirty individuals participated in Experiment 1, 30 in Experiment 2, 31 in Experiment 3 and 32 in Experiment 4. All participants were non-musicians and scored an average of 17.45 (SD = ± 7.60, range: 7 - 39) on the musical training subscale of the Goldsmiths Musical Sophistication Index (Gold-MSI) questionnaire[73]. All participants reported normal or corrected-to-normal vision and an absence of neurological or psychiatric diseases. They gave written informed consent prior to experiment and received a monetary compensation of ¥ 50 per hour for their participation. The study was approved by the Ethics Committee of the Institute of Psychology, Chinese Academy of Sciences.

### Stimuli

The stimuli comprised ten musical pieces selected from the 371 four-part chorales by Johann Sebastian Bach (Breitkopf Edition, Nr. 8610). The original musical scores were checked manually to ensure inclusion of only quarter notes, eighth notes and sixteenth notes, while removing ties across notes to facilitate repetition. Importantly, we eliminated fermatas from the original musical pieces to prompt participants to parse the musical pieces by structural cues rather than rhythmical or acoustic cues for phrasal structures (such as pauses between phrases and lengthened notes). The duration of the musical pieces ranged from 22.59 s to 59.30 s (39.82 ± 11.49 s, mean ± SD).

In Experiment 1, we created a reverse version by completely reversing the order of beats in each original piece (for an example piece, see Fig 1B). Therefore, the basic musical contents were equal in both versions, but the harmonic progressions were manipulated in the reverse version which would affect listeners’ expectation or familiarity of the musical pieces.

For the rhythm condition in Experiment 2, the stimuli consisted of frequent standard tones (440 Hz) with timing structures identical to the original pieces. To ensure participants’ attention remained focused, we randomly inserted one to three chords deviating in timbre (guitar) into two additional pieces selected from the musical stimuli. These two pieces containing timbre deviants were excluded from further analyses.

In Experiment 3, we extracted twenty-four melodic sequences, each comprising two phrases, from the original pieces used in Experiment 1. Each phrase consisted of eight chords. A pitch deviant note occurred randomly within the second phrase of each melody with equal probability. To adjust difficulty levels, we created pitch deviants by shifting the melody in the upper two voice parts (soprano and alto) upwards by one to two octaves. Each sequence was manipulated into two versions with the deviant inserted in different position.

Experiment 4 utilized visual stimuli, with the target being a red circle with a diameter of 50 pixels (RGB: 255, 0, 0), displayed at the center on a black background (RGB: 0, 0, 0).

All stimuli were created and exported as wav files with a piano timbre using the MuseScore 3 software. In Experiments 1, 3 and 4, musical stimuli were played at a tempo of 85 beats per minute, while Experiment 2 employed three tempi (66, 85 and 120 beats per minute). Therefore, the note rate for each musical piece at a tempo of 85 beats per minute was around 2.833 Hz, the beat rate was around 1.416 Hz, and the phrase rate was around 0.177 Hz. Musical pieces were presented at a sampling rate of 44100 Hz, and the intensity level of each piece was normalized to 70 dB SPL.

### Experimental design and procedure

Experiment 1: The twenty musical pieces (10 pieces × 2 versions) were presented in 4 blocks, each containing 5 pieces. In each block, participants listened to each of the 5 pieces three times in a randomized order. Throughout the experiment, participants were required to fixate on a white fixation cross at the center of a black screen. After the third presentation of each musical piece, participants pressed keys to rate how much they liked it using a 5-point Likert scale, with 1 indicating low preference and 5 indicating high preference. After providing their rating, the next musical piece commenced after a delay of 3∼3.5 s (see Fig 1A). Each block lasted around 10 minutes and participants allowed to rest between blocks.

Experiment 2: The melody condition and the rhythm condition were presented in separate blocks, with the conditions alternating every 5 trials. Stimuli were presented to participants according to a balanced Latin square design for every condition, and the order of the two conditions was counterbalanced across participants. Before the main experiment, participants underwent a 3-min rest period while looking at a blank screen to obtain a baseline. During each block, participants were instructed to focus on the fixation cross displayed on the screen and press the spacebar as soon as possible upon detecting a timbre deviant. Participants received training until they achieved an accuracy level of 80%.

Experiment 3: Participants were presented with two versions of the twenty-four melodic sequences, each presented twice in a pseudorandomized order, ensuring that no sequence or version was repeated consecutively. Participants were asked to carefully listen to the sequences and press the spacebar as soon as they detected a pitch deviant, while maintaining fixation on the central cross on the screen. Before the experiment, participants underwent a training session, concluding once they achieved an accuracy score above 80% on the detection task.

Experiment 4: Participants engaged in a visual target detection task while ten musical pieces, used as the original version in Experiment 1, were presented twice in a pseudorandomized order. Concurrently, the fixation cross at the center of the screen occasionally transformed into a red circle. The visual target appeared either synchronously with the beat onset (on-beat condition) or randomly at one of nine temporal positions between two beats (off-beat condition). It was ensured that 1) each musical piece contained at least one target and up to three for each condition with the number of targets per condition was the same, and 2) no piece was presented twice consecutively with visual targets occurring at different positions for the same piece. Participants were instructed to observe the fixation cross while the auditory stimuli played, and press the spacebar as soon as possible once sighting the red circle on the screen. A training session was given before the experiment until participants achieved a target detection accuracy exceeding 80%.

At the end of each experiment, participants completed the Gold-MSI questionnaire and MET to measure their musical ability. The English version of the Gold-MSI questionnaire was translated into Chinese by the experimenters. Participants in Experiment 4 did not complete the MET due to the duration of the task.

### Data acquisition

Eye-tracking data were recorded in all experiments using an Eyelink Portable Duo system (SR Research, Ontario, Canada), with a sampling rate of 500 Hz. Participants maintained a consistent eye-to-monitor distance of 95 cm, with the eye tracker positioned approximately 60 cm from their eyes. Before each experiment, a nine-point standard calibration and validation test was performed. Then a fixation cross appeared on the monitor with a resolution of 1,920 × 1,080 pixels. A re-calibration procedure was applied after each break to ensure accuracy and consistency.

In Experiment 1, EEG data were recorded using a 64-channel Quik-Cap based on the international 10–20 system (Neuroscan, VA, USA), with a sampling rate of 1000 Hz. The EEG was referenced to the average of all electrodes, with skin/electrode impedance maintained below 5 kΩ . Additionally, four electrodes were used to record electrooculography (EOG): two electrodes were placed at the outer canthus of each eye, while another two electrodes were placed above and below the left eye to record the horizontal and vertical EOG signals, respectively. Additionally, DWI data were collected from the same cohort using a 3.0 T MRI system (Simens Magnetom Prisma) with a 20-channel head coil with following parameters: repetition time (TR) = 4000 ms, echo time (TE) = 79 ms, field of view (FOV) = 192 × 192 mm^2^, voxel size = 1.5 × 1.5 × 1.5 mm^3^, diffusion-weighted gradient directions = 64, b-values = 1000 s/mm^2^ and 2000 s/mm^2^, b0 non-weighted images = 5.

### Data preprocessing

Preprocessing of eye-tracking and EEG data were performed in MATLAB (The MathWorks, Inc., Natick, MA) using the Fieldtrip toolbox[74] and custom scripts.

For eye-tracking data, blink detection was the primary event of interest. Blinks were identified by the Eyelink tracker’s on-line parser, which flagged instances where the pupil in the camera image was absent or severely distorted by eyelid occlusion. The time series were then divided into epochs corresponding to the duration of each musical or pure-tone sequence, aligning with the experimental trials. In Experiment 2, baseline data was derived from three continuous trials, each lasting 60 s. Within each trial, data were represented as binary strings, with values set to 1 during blink occurrences and 0 otherwise. Blinks shorter than 50 ms or longer than 500 ms were excluded from the analysis. Subsequently, mean blink duration and blink rate were calculated for each participant. Blinks with mean durations exceeding three standard deviations from the data’s mean, and trials with blink rates exceeding three times the standard deviation, were removed. Following preprocessing, eye-tracking data were down-sampled to 100 Hz for further analysis.

For continuous EEG data, single trials were extracted spanning 3 s before stimulus onset to 3 s after stimulus offset. All data were band-pass filtered between 0.7 and 35 Hz using a phase-preserving two-pass fourth-order Butterworth filter, supplemented with a 48- to 52-Hz band-stop filter to eliminate line noise. Then the data were down-sampled to 100 Hz to match the sampling rate of the eye-tracking data. An independent component analysis[75] with 30 principal components, implemented in Fieldtrip, was performed to remove EOG and electrocardiography (ECG) artifacts. Finally, we concatenated all trials into one matrix to derive the most common component of the EEG signals reflecting neural responses to musical pieces, using MCCA[76]. The first MCCA component was projected back to each participant and each trial, and used for subsequent analysis.

For DWI data, MRtrix3 and FSL software[77, 78] were employed to preprocess the raw data. Preprocessing steps included denoising, reducing the ringing artifacts, eddy current and motion correction, and bias field correction using the N4 algorithm provided in Advanced Normalization Tools. Gradient directions were also corrected after eddy current and motion correction.

### Music amplitude modulation spectrum

To derive the amplitude envelope of the musical stimuli in Experiment 1, we constructed 64 logarithmically spaced cochlear bands between 50 to 4000 Hz using a Gammatone filterbank[79]. The amplitude envelope of each band was then extracted from the acoustic waveform using the Hilbert transform and down-sampled to 100 Hz. For each musical piece, the amplitude envelopes of the 64 bands were averaged, and the modulation spectrum of each piece was calculated using the FFT with zero-padding of 8000 points, followed by taking the absolute value of each frequency point. Next, we normalized the modulation spectrum of each piece by dividing it by the norm of its modulation spectrum. Lastly, we estimated the mean amplitude modulation spectrum across the 10 pieces for each condition and showed them in Fig 1C.

### Blink and neural synchronization

To access whether blink and neural signals could synchronize to musical structures, we used a frequency domain analysis. The analysis window for blink data varied depending on the task. In summary, we analyzed the time series from 1 s after stimulus onset to 1 s before stimulus offset in Experiments 1 and 2, and we selected the time series from 1 s after stimulus onset to the onset of the first target in Experiments 3 and 4. The blink data for each trial and participant were then transformed into the frequency domain using a Fast Fourier Transform with zero-padding of 6000 points. Given the variable trial lengths, we averaged the amplitude spectra across the 10 pieces for each version and repetition in the frequency domain. Next, we identified the lower and upper bounds on the significant frequency ranges among all independent variables showing robust blink and neural tracking of musical structures. We then averaged the amplitude spectra within the frequency range for each variable for further investigation. For the baseline in Experiment 2, we first averaged the data across three trials in the temporal domain and then applied the same procedure for conversion to the frequency domain.

### Mutual information

In the above frequency-domain analysis, we observed robust blink synchronization to musical sequences primarily centered around the beat rate. To estimate the statistical dependency between the neural response and the eye-tracking data, we focused on the EEG power at the beat rate and computed mutual information (MI) between it and the blink signal. First, we extracted the time-frequency distribution of EEG data at the beat rate using a Morlet wavelet (with a sliding window length of 3) implemented in Fieldtrip, with steps of 10 ms. Next, we converted the output, corresponding to the EEG power, to a decibel scale using a pre-stimulus baseline ranging from -1500 to - 500 ms. Then, MI between the blink signal and EEG power was calculated using Gaussian Copula Mutual Information[80] at different lags: the blink signal was shifted against the EEG power from -200 to 200ms in 20-ms steps. Finally, we summed the MI values across lags and averaged them across 10 pieces for each version and repetition for further analysis.

### Temporal response function

What’s the relationship between ocular activity and neural response in the time domain? To reveal the temporal dynamics of the neural response to blink activity, we employed the mTRF toolbox[81] to estimate the temporal response function (TRF) of EEG power at the beat rate, utilizing the blink onset vector as the input stimulus feature. The TRF, denoted as w, was estimated using the following equation:

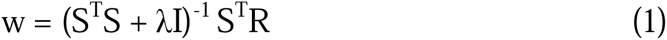

Here, S is the lagged time series of the input stimulus feature (the blink onset vector obtained from the eye-tracking data), R is the continuous neural response at the beat rate, I is the identity matrix, and λ is a constant ridge parameter.

The TRF for each musical piece and participant was calculated using the blink onset vector and EEG power from 1 s after stimulus onset to 1 s before stimulus offset. The time lags ranged from 2500 ms before blink onset to 2500 ms after, with λ set to 0. Then we averaged the TRF weights across 10 pieces for each version and repetition for further permutation test (described in the following section), and extracted the TRF weights at the peak time points for all independent variables for further comparison.

### Fiber tractography

Fiber tractography based on high angular resolution diffusion imaging (HARDI) was performed using MRtrix3. Firstly, 3-tissue (white matter, gray matter, and cerebrospinal fluid) response functions were obtained by the command ‘dwi2response dhollander’[82]. Secondly, fiber orientation distributions (FOD) of each voxel were estimated using multi-shell multi-tissue constrained spherical deconvolution algorithm[83]. Thirdly, the whole-brain probabilistic tractography was performed to generate ten million streamlines for each participant based on iFOD2 tracking algorithm[84]. Lastly, track filtering was performed to attain 1 million streamlines[85].

The target segments of the AF and SLF were defined according to a simple classification system[86]: the dorsal SLF (dSLF), namely the classical SLF II, connects the Geschwind’s area and the dorsolateral frontal area; the ventral SLF (vSLF) corresponds to the SLF III or anterior AF connecting the Broca’s area and the Geschwind’s area; the posterior SLF (pSLF) corresponding to the SLF-tp or posterior AF connects the Geschwind’s area and the Wernicke’s area; the AF, alternatively known as the classical AF or long AF, connects the Broca’s area and the Wernicke’s area. Four ROIs (Geschwind’s area: super marginal gyrus and angular gyrus; dorsolateral frontal area: posterior superior frontal gyrus and middle frontal gyrus; Broca’s area: opercular part of inferior frontal gyrus and inferior part of precentral gyrus; Wernicke’s area: posterior superior temporal gyrus and middle temporal gyrus) were extracted from individual anatomical image parceled by Freesurfer for each hemisphere to dissect bilateral segments of AF/SLF. S6 Fig shows fiber tractographies of the AF and SLF in bilateral hemispheres from one participant.

We extracted three indexes to measure the microstructural properties of fibers: FA derived from DTI as well as NDI and ODI derived from the NODDI model[55]. The mean FA, NDI and ODI values of each segment were extracted for each participant. LI for each parameter was subsequently calculated using the following equation:

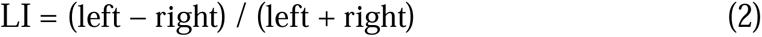

### Behavioural measure

Accuracy and reaction time were recorded in Experiments 3 and 4. Detection accuracy was defined as the ratio of correctly detected targets within one beat duration to the total number of targets presented. Only correct trials were included in the reaction time analysis.

### Blink rate

In this analysis, we derived blink time series within a window from -0.353 to 0.353 s in relative to the beat onset for each beat, determined by the tempo (85 beats per minute). The blink rate at each time point was defined as the average value across trials for each participant. Results are illustrated in S5 Fig.

### Statistical analysis

All statistical analyses were calculated using MATLAB and SPSS 20.0 (IBM Corp., Armonk, N.Y., USA). Group statistical analyses were performed on the data of all participants. Prior to analysis, outliers were identified and removed using the mean detection method.

For behavioural data, statistical differences between conditions (Figs 1D and 6E) were evaluated using a paired t-test with a threshold of *p* = 0.05. For the ocular and neural activities, all statistical analyses (Figs 2B, 3B, 3E, 5D left and S2B) were performed with repeated-measures ANOVA to examine the effects of reversal manipulation, stimulus repetition, stimulus type and tempo. To address the issue of multiple comparisons, the statistical significance level was set at a corrected *p* < 0.05 using the Bonferroni correction method. The Greenhouse-Geisser correction was used when the assumption of sphericity was violated. The individual differences in blink response amplitude (Fig 4A) were evaluated by a median split approach. Statistical differences between groups (Fig 4B and 4D) were evaluated using an independent t-test with a threshold of *p* = 0.05. For neural activities, group comparisons (Fig 4C) were evaluated using a Mann-Whitney U test. All statistical tests performed were two-tailed. Correlations between blink/neural responses and behavioural indexes (Figs 2C, 2E, 5D right, 6C and S2C) were estimated using Spearman’s rank correlation coefficients or Pearson’s correlation coefficients according to the data distribution.

To identify the significant frequency ranges showing blink/neural tracking of auditory sequences (Figs 2A, 2D, 5C, 6B and 6D), we completed a surrogate test. Blink/neural data underwent the same Fourier transform processing steps described above, yielding a complex number at each frequency point for each trial and participant. Then the data were phase shuffled in the frequency domain by multiplying the data with a complex number which has a magnitude of 1 and a randomly generated phase between 0 and 2π. As for the frequency domain analysis, we averaged the amplitude spectra of surrogate data across trials for each condition. This procedure was repeated 1000 times, creating a null distribution of 1000 surrogate amplitude modulation spectra for each condition, from which the 99^th^ percentile was chosen as the significant threshold. Finally, we z-scored the raw modulation spectra with respect to the mean of the surrogate distribution for each condition. For baseline data (Fig 5B), the surrogate test procedure was identical, except for data surrogation in the temporal domain.

To assess the statistical significance of the coupling between the blink signal and the beat-rate EEG response (Fig 3A), we conducted a permutation test. Specifically, the beat-rate EEG power for each participant and trial was circularly shifted by a random shift amount within the time series, and MI between the time-shifted data and blink signal was recalculated. This procedure was repeated 1000 times to create a null distribution of surrogate MI values for each version and repetition, from which a p-value was obtained by counting how many surrogate MI values exceeding the empirical MI value.

To confirm that the rise in TRF weights before blink onset was attributable to pre-onset prediction for the beat-rate neural response to blink (Fig 3D), a permutation test was completed following the same procedure used for Fig 2D. The TRF was estimated between the new blink signal and EEG power 1000 times, and a threshold of a one-sided alpha level of 0.01 was derived for statistical significance.

To assess the significantly different frequency ranges between two groups (S4 Fig), we conducted a cluster-based permutation test. Firstly, we compared blink amplitude of two groups with a two-tailed independent t-test. Selected frequency points where t-values exceeded a threshold of 0.05 were clustered in connected sets on the basis of spectral adjacency and the t-values within every cluster were summed. The maximum cluster sum was then compared to the maximum cluster sums that were obtained after randomly swapping the label of the blink amplitude between the two groups 1000 times. A p-value was derived as the ratio of maximum clusters that exceeded the real maximum cluster.

## Acknowledgments

We thank Lidongsheng Xing and Jingwen Wang for their assistance with data collection; Pauline Larrouy-Maestri and Hou Chen for their assistance in preparing music materials; Baishen Liang, Xiang Li, Lei Zhang, Lyu Baihan and Peiqing Jin for their help with data processing. We thank Robert Zatorre for his comments on a previous version of the manuscript.

## Supporting information

**S1 Fig.**
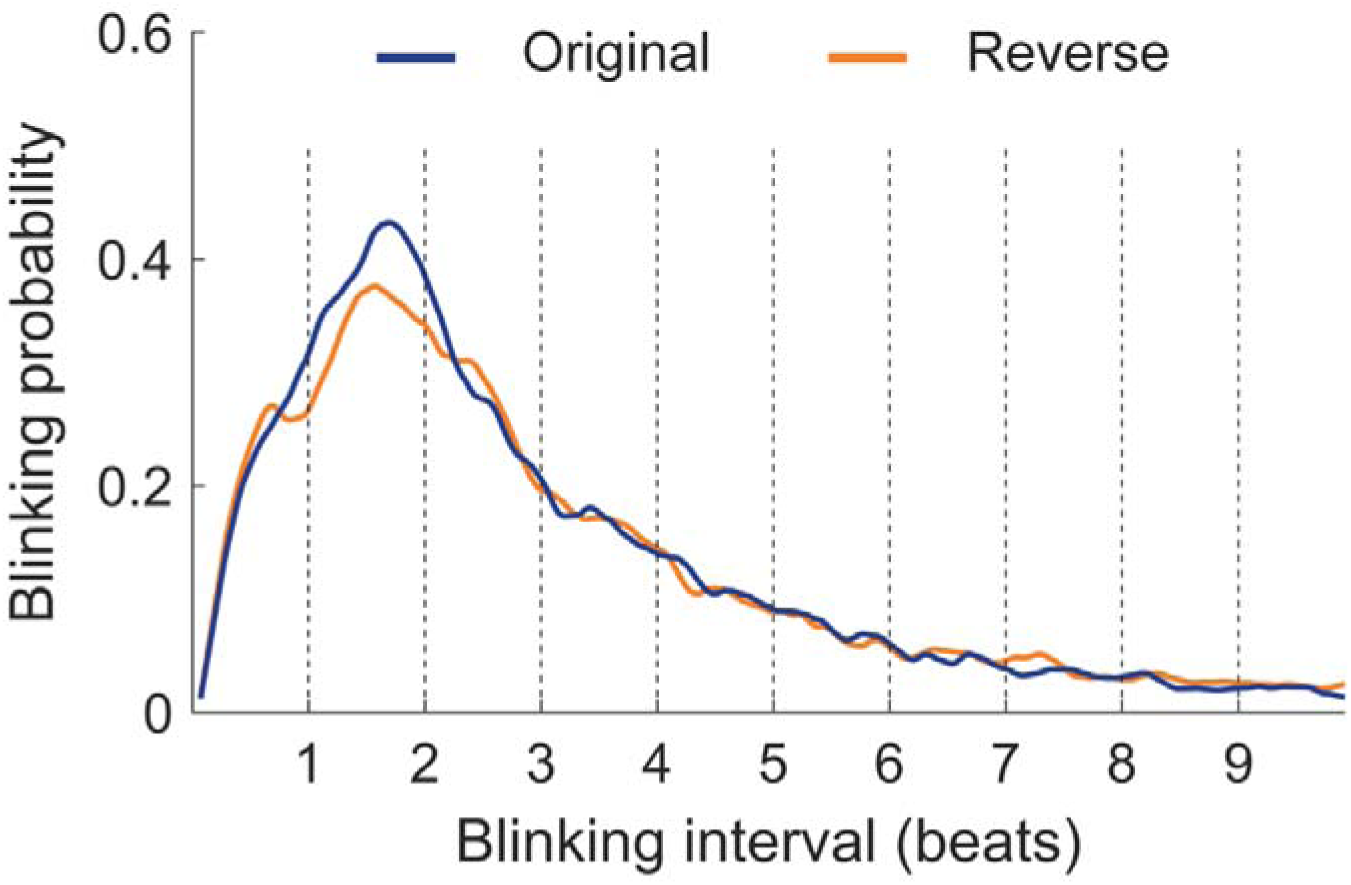
Blink probability at different time interval for two versions. The length of each musical beat is 0.706 s. The figure displays that the inter-blink interval is most often one to two beats, consistent with what we observe in Fig 2.

**S2 Fig.**
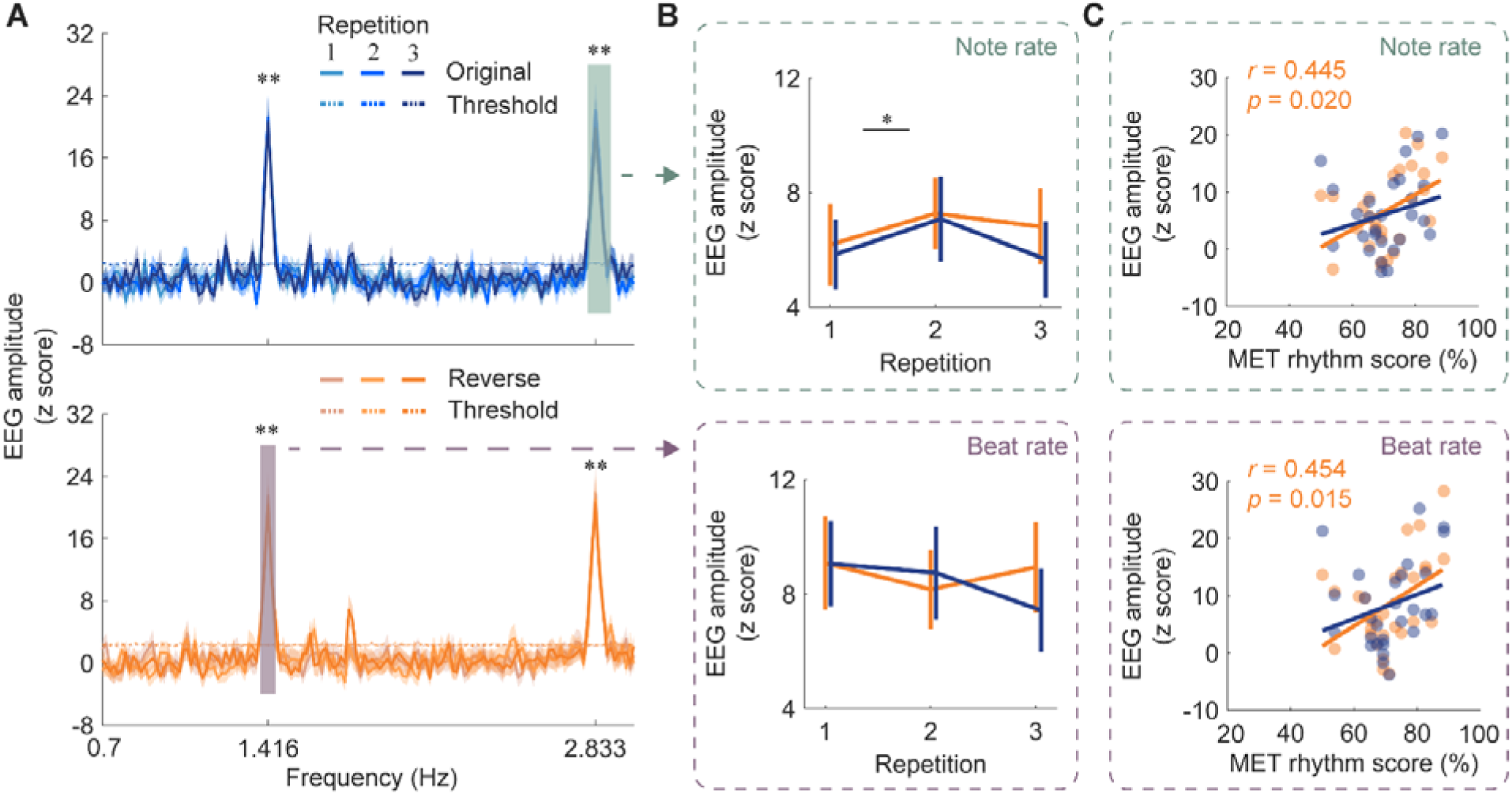
Neural entrainment to musical beats and notes. **(A)** Neural response spectra for two versions consistent with Fig 2D. The colored boxes indicate the frequency ranges where the amplitude was above the threshold. **(B)** EEG amplitude at note and beat rates. Error bars denote 1 SEM across participants. Note-rate EEG response increased with repeated exposure of music. **(C)** Correlation between the MET rhythm score and EEG amplitude at note and beat rates. The two EEG responses were both correlated with the MET rhythm score for the reverse version. Colored dots indicate individuals. * *p* < 0.05, ** *p* < 0.01.

**S3 Fig.**
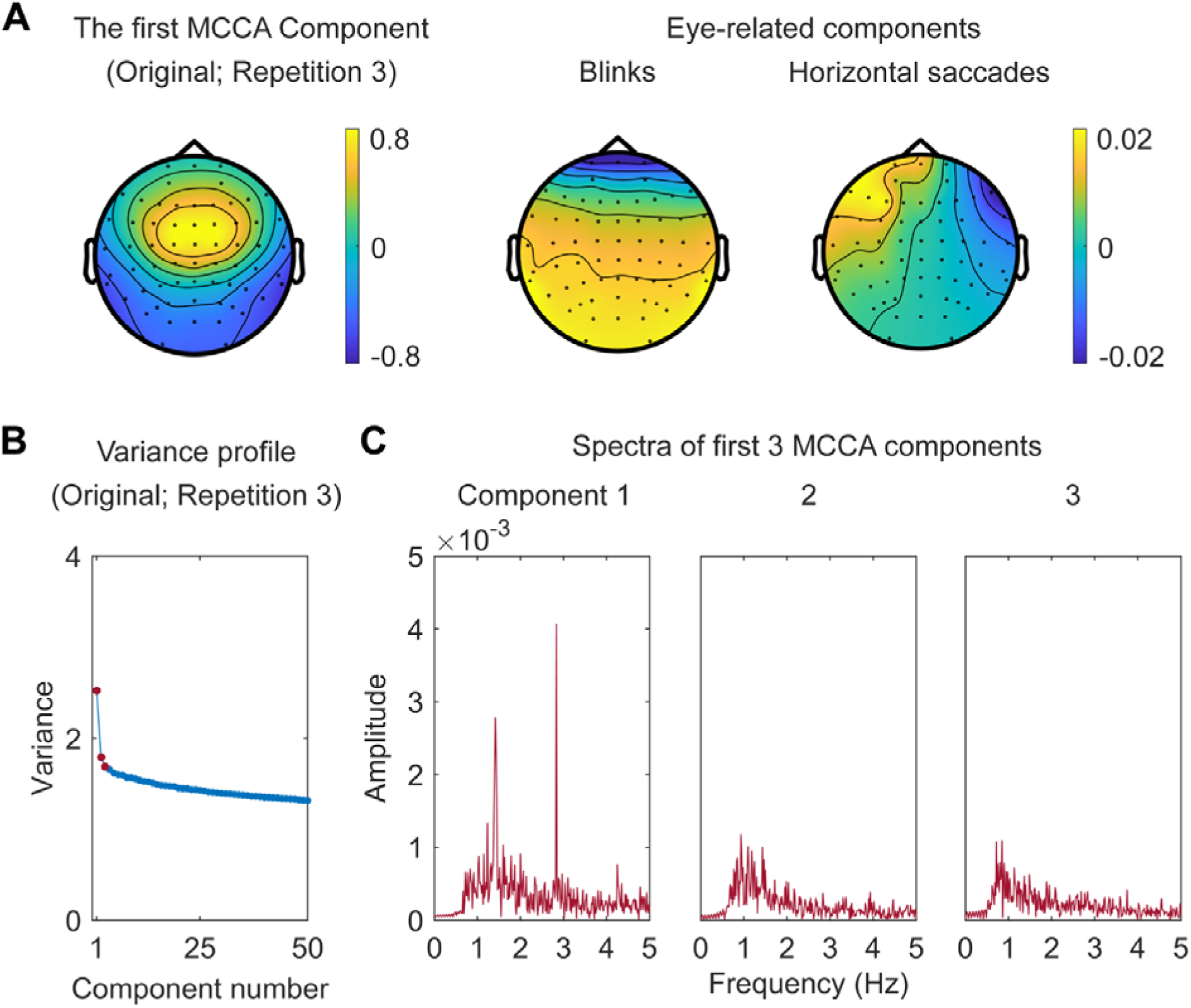
Multiway canonical correlation analysis (MCCA). **(A)** Scalp topographies for the first MCCA component for the third presentation in the original condition (left) and eye-related components extracted using independent component analysis (ICA) (right). **(B)** Variance of summary components (SCs) extracted from MCCA. The variance of the first SC explains a considerable amount of variance. **(C)** Amplitude spectra of first three MCCA components for the third presentation in the original condition. The spectrum of the first component shows amplitude peaks corresponding to the beat rate and note rate (the first harmonic of the beat rate), further supporting the first MCCA component, but not other components, contained the neural signals induced by beat and note structures in the music pieces. Similar results are shown for other repetitions and other reversal conditions.

**S4 Fig.**
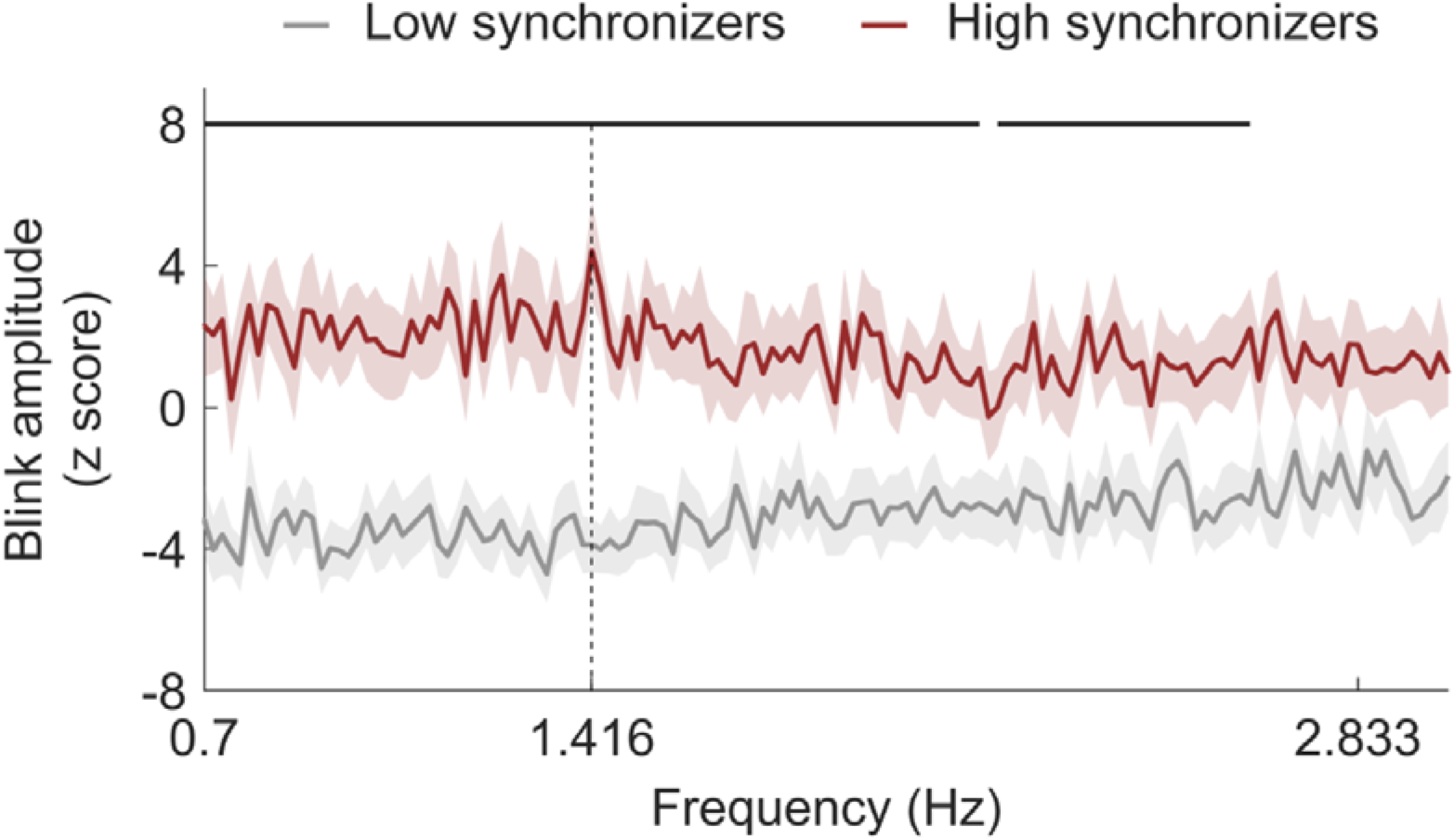
The blink amplitude spectrum for two groups. Difference in blink amplitude between the groups (low synchronizers: gray, high synchronizers: red). Horizontal black lines at the top mark significant difference (cluster-based permutation test, *p* < 0.01).

**S5 Fig.**
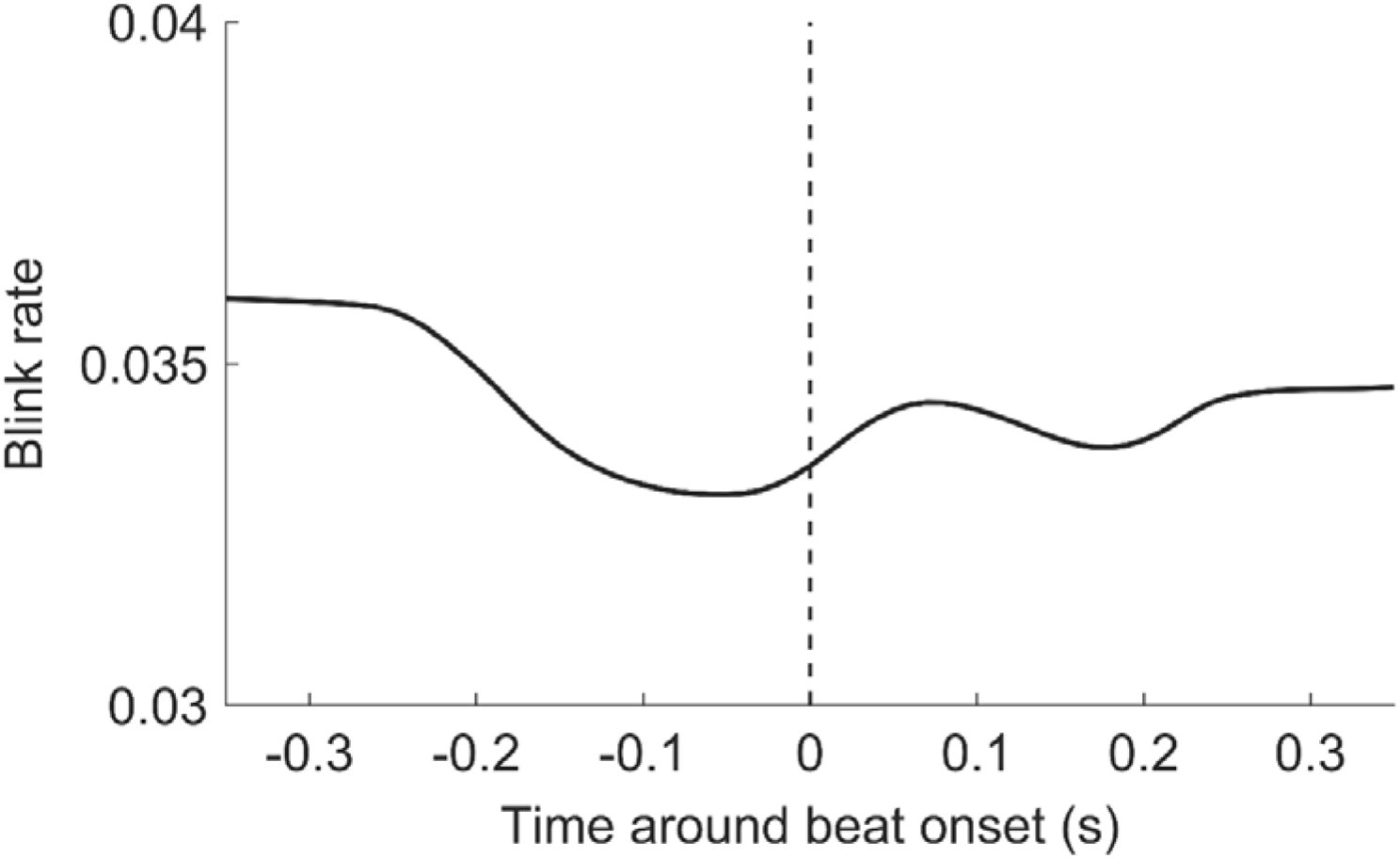
The time series of blink rate in Experiment 4. Blinks were more likely to occur around the beat onset. The vertical dashed line represents beat onset. N = 32 participants.

**S6 Fig.**
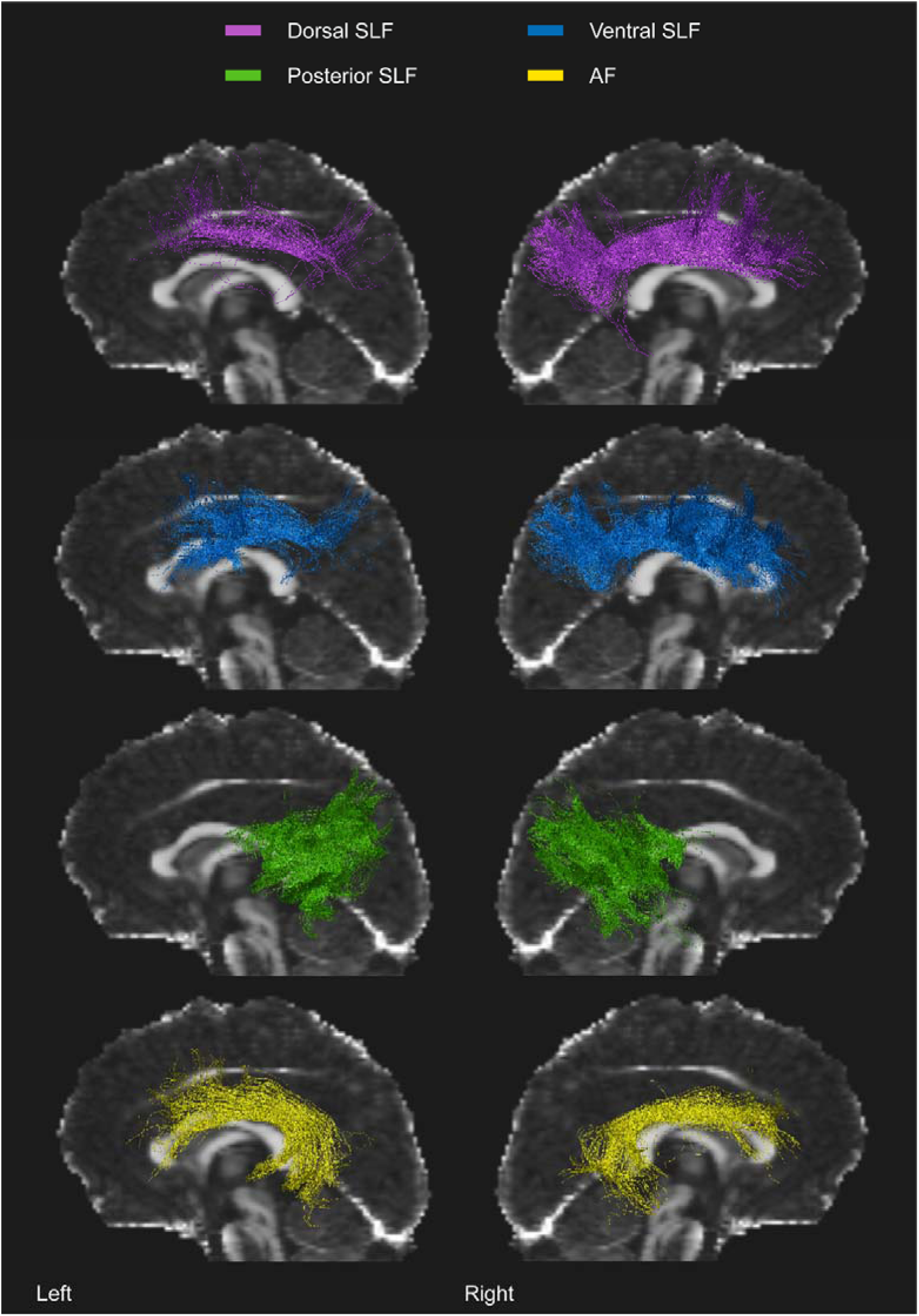
The tractography of AF and SLF in bilateral hemispheres from a participant. Dorsal, ventral and posterior segments of SLF are depicted in purple, blue and green. The AF is depicted in yellow.

**S1 Table.** Group comparisons of microstructural and laterality index between low synchronizers and high synchronizers. Values are *t* (*p*). P was estimated by two-sided independent t-test with a threshold of *p* = 0.05. * *p* < 0.05.

|  | Microstructural index |  |  | Laterality index |  |  |
| --- | --- | --- | --- | --- | --- | --- |
|  | FA | NDI | ODI | FA | NDI | ODI |
| L dorsal SLF | -1.257<br>(0.221) | 0.585<br>(0.564) | 2.025<br>(0.054) | -0.877<br>(0.390) | -0.004<br>(0.997) | 0.788<br>(0.438) |
| R dorsal SLF | -0.323<br>(0.749) | 0.539<br>(0.595) | 1.286<br>(0.211) |  |  |  |
| L ventral SLF | -0.181<br>(0.858) | 1.413<br>(0.170) | 0.995<br>(0.330) | 0.014<br>(0.989) | 1.629<br>(0.116) | 0.195<br>(0.847) |
| R ventral SLF | -0.424<br>(0.675) | 0.159<br>(0.875) | 1.576<br>(0.128) |  |  |  |
| L posterior SLF | 1.050<br>(0.304) | 1.929<br>(0.066) | -0.190<br>(0.851) | 1.778<br>(0.088) | 0.863<br>(0.397) | -1.452<br>(0.160) |
| R posterior SLF | -0.322<br>(0.750) | 0.964<br>(0.345) | 1.141<br>(0.265) |  |  |  |
| L AF | -0.097<br>(0.924) | 1.830<br>(0.080) | **2.272**<br>**(0.032) \*** | 0.095<br>(0.925) | 0.461<br>(0.649) | -0.076<br>(0.940) |
| R AF | -0.109<br>(0.914) | 1.157<br>(0.259) | 1.446<br>(0.161) |  |  |  |

